# Transcription start site profiling uncovers divergent transcription and enhancer-associated RNAs in *Drosophila melanogaster*

**DOI:** 10.1101/165639

**Authors:** Michael P. Meers, Karen Adelman, Robert J. Duronio, Brian D. Strahl, Daniel J. McKay, A. Gregory Matera

## Abstract

**Background:** High-resolution transcription start site (TSS) mapping in D. melanogaster embryos and cell lines has revealed a rich and detailed landscape of both cis- and trans-regulatory elements and factors. However, TSS profiling has not been investigated in an orthogonal in vivo setting. Here, we present a comprehensive dataset that links TSS dynamics with nucleosome occupancy and gene expression in the wandering third instar larva, a developmental stage characterized by large-scale shifts in transcriptional programs in preparation for metamorphosis.

**Results:** The data recapitulate major regulatory classes of TSSs, based on peak width, promoter-proximal polymerase pausing, and cis-regulatory element density. We confirm the paucity of divergent transcription units in D. melanogaster, but also identify notable exceptions. Furthermore, we identify thousands of novel initiation events occurring at unannotated TSSs that can be classified into functional categories by their local density of histone modifications. Interestingly, a sub-class of these unannotated TSSs overlaps with functionally validated enhancer elements, consistent with a regulatory role for “enhancer RNAs” (eRNAs) in defining transcriptional programs that are important for animal development.

**Conclusions:** High-depth TSS mapping is a powerful strategy for identifying and characterizing low-abundance and/or low-stability RNAs. Global analysis of transcription initiation patterns in a developing organism reveals a vast number of novel initiation events that identify potential eRNAs as well as other non-coding transcripts critical for animal development.

## Introduction

Transcription initiation constitutes the first step in gene expression, and thus its fidelity is of utmost importance for proper regulation of gene expression (Sainsbury *et al.* 2015). Initiation begins when the pre-initiation complex (PIC) assembles on exposed DNA at a promoter region upstream of the transcription start site (TSS) [1]. Through the action of both active and passive mechanisms, promoters are disproportionately depleted of nucleosomes, and are thus available for PIC assembly. These mechanisms include binding of specialized transcription factors [2], activity of nucleosome remodelers [3], and sequence-dependent likelihood of nucleosome assembly [4,5]. The interplay of these factors is important for generating transcripts with temporal and spatial specificity [6], and for suppressing initiation from cryptic or developmentally inappropriate sites that may otherwise be competent for initiation [7,8]. Regulation of initiation has important implications for cell differentiation, where activation of developmentally significant “master regulator” genes can alter gene expression regimes that define cellular morphology and identity. For instance, the expression of a handful of transcription factors associated with pluripotency is sufficient to transform differentiated cells into induced pluripotent stem cells [9].

Transcription initiation can be regulated at several levels. Prior to RNA polymerase II (RNA pol II) engaging with DNA at the TSS, nucleosome depletion directly upstream of the TSS facilitates assembly of the PIC and other general transcription factors. This stereotypical ‘minus-1’ nucleosome depleted region (NDR) is conserved across eukaryotes [10,11], and is highly correlated with transcription initiation activity. Factors that alter the likelihood that a NDR occurs will also alter the propensity of RNA pol II to initiate at that site. Similarly, transcription factor binding to *cis* elements in the promoter results in displacement of nucleosomes. Additional descriptive characteristics of transcription initiation activity, such as the breadth or distribution of initiating polymerases across a given domain [12,13], correlate with gene expression outcomes. However, it is not known whether these factors play a role in proper regulation of gene expression.

Furthermore, transcription initiation has been shown to occur in divergent directions, with unclear consequences for gene expression [14,15]. In most cases transcripts that are produced in the antisense direction relative to an annotated gene are rapidly degraded [16]. Divergent transcription initiation is a common feature in mammals [17], and is observed across annotated TSSs and enhancer regions [18]. However, it is still unclear whether bidirectional transcription is functionally relevant to gene expression, particularly because certain cell types, including *D. melanogaster* S2 cells, appear to be largely devoid of divergent initiation [19].

A final initiation-related regulatory step occurs after PIC assembly, when RNA pol II transcribes ~50-100nt into the gene body before it is subject to promoter proximal pausing. Pausing can act as a regulatory step to help integrate signals or it can prepare promoters for rapid activation [20,21]. Although the dynamics of polymerase pausing are well understood in cell culture [19–21], to date there have been few studies that have comprehensively characterized pausing *in vivo* [22].

At potential sites of transcription initiation outside of annotated TSSs, in most cases surveillance and degradation by the nuclear exosome occurs rapidly [23,24]. This degradation is likely important because initiation at non-canonical or cryptic promoters can interfere with coding transcripts or create a deleterious load of non-functional ones, including dsRNAs [25]. In general, sites of initiation unassociated with annotated gene promoters have a high propensity for nucleosome occupancy, and are energetically unfavorable for assembly of the PIC [4,5]. However, in the budding yeast *Saccharomyces cerevisiae,* genetic perturbations that cause cryptic initiation in coding regions are tolerated [7,8,26]. Furthermore, there is evidence that transcription from unannotated promoters may also serve beneficial functions [27], particularly at enhancer regions [28], which have been shown to produce enhancer RNAs (eRNAs) that may play regulatory roles [14,15,17]. Whereas cryptic and unnanotated transcription has been extensively characterized and described in *S. cerevisiae* (e.g. [29]), it is less well characterized in metazoans.

Here, we present a detailed characterization of matched Start-seq [19], ATAC-seq [30], and nuclear RNA-seq datasets in *D. melanogaster* 3^rd^ instar larvae [31]. From these data, we were able to annotate larval TSSs with nucleotide resolution, and analyze connections between local cis-regulatory motifs, TSS shape, pausing activity, and divergent transcription. Additionally, we identified thousands of unannotated initiation events, and used existing datasets for histone post-translational modifications (PTMs) and validated enhancer regions to impute their functions. Our findings are among the first to detail the global initiation patterns in a developing organism, uncovering a vast number of new initiation events that define likely enhancer RNAs and transcripts critical for animal development.

## Results

### Start-seq signal correlates with nucleosome depletion, gene expression, and promoter proximal pausing

To characterize the genome-wide landscape of gene expression, transcription initiation, and chromatin accessibility in third instar *Drosophila melanogaster* larvae, we carried out rRNA-depleted total nuclear RNA-seq, Start-seq, which quantifies short, capped, nascent RNAs that represent newly initiated species [19,21], and ATAC-seq, which quantifies transposase-accessible open chromatin [30], as previously described [31]. For every annotated gene, we assigned the dominant Start-seq peak most likely to represent its bona-fide TSS from its most frequently used start site in order to cross-compare open chromatin, initiation, and gene expression values within each gene (Fig. 1A). As shown in Figure 1B, ATAC-seq signal is highest in the 150nt upstream and 50nt downstream of the TSS, corresponding to the expected location of a promoter-proximal NDR [10]. Additionally, Start-seq signal accumulates robustly and almost exclusively within the ~50nt directly downstream of the assigned TSSs (Fig. 1B), consistent with expected signal distributions from previously reported Start-seq analyses [19]. Importantly, the first nucleotide in the 5’ read of each Start-seq read pair acts as a proxy for the first transcribed nucleotide in the nascent mRNA chain [19], enabling bona-fide TSS mapping at single base-pair resolution.

**Figure 1:**
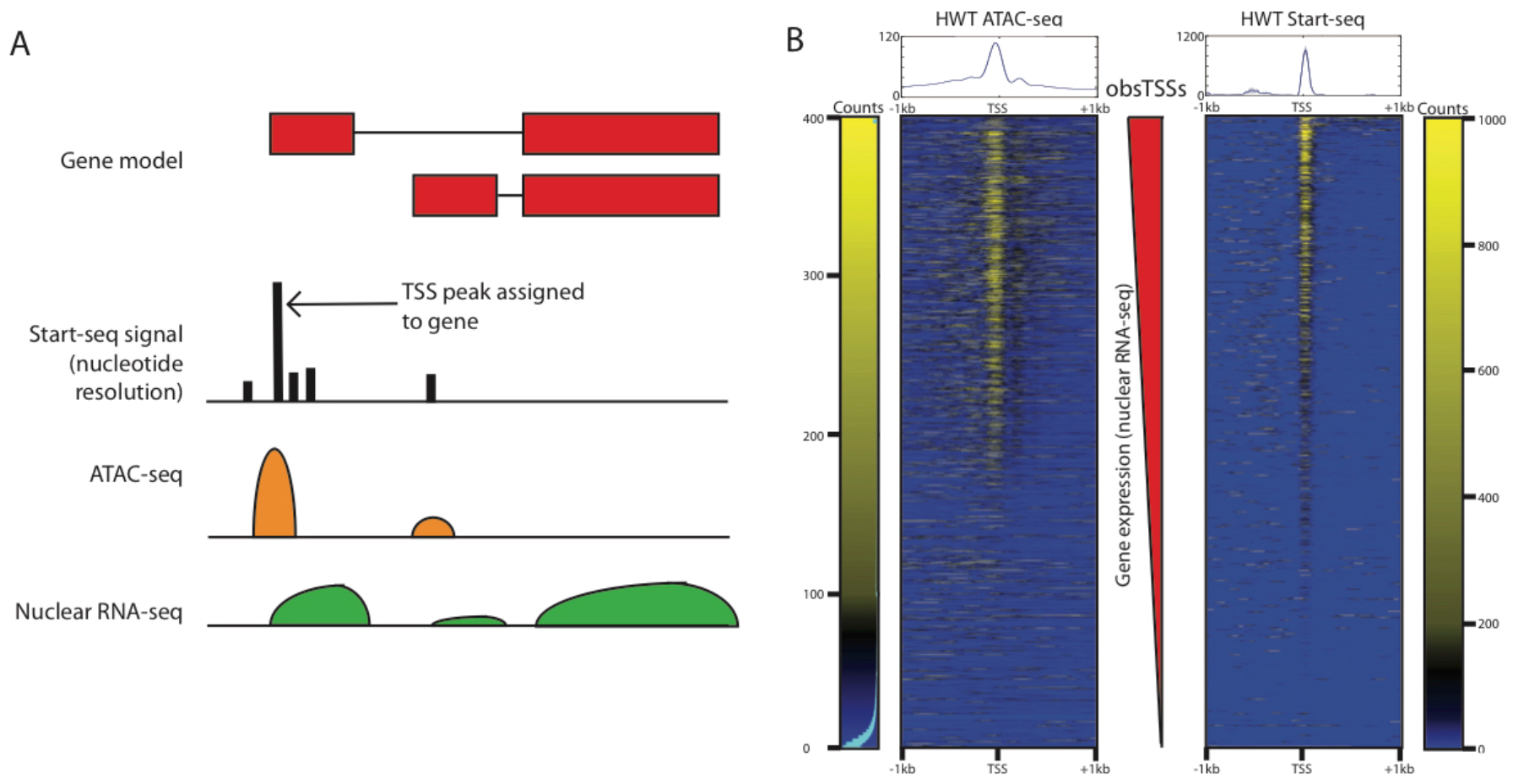
Comparison of ATAC-seq and Start-seq data. A) Schematic describing assignment and linkage of Start-seq, ATAC-seq, and nuclear RNA-seq within a single gene. B) Heatmap for ATAC-seq (left) and Start-seq (right) signal mapping at annotated transcription start sites (obsTSSs), ordered by increasing nuclear RNA-seq signal.

Nucleosomes are barriers to transcription factor binding and PIC assembly [11]. Accordingly, the extent of chromatin accessibility has been shown to correlate with the level of gene expression [10,11], and thus should correlate well with the level of transcription initiation. To evaluate these expected relationships on a gene-specific level, we used the most frequently used start site for each gene in the genome to assign a discrete value for chromatin accessibility, transcription initiation, and nuclear RNA-seq gene expression level, and performed correlation comparisons between each pair of values across all genes. Although Start-seq intensity generally correlates well with overall gene expression, ATAC-seq levels correlate poorly with Start-seq (Fig. S1A). Curiously, both Start-seq and ATAC-seq correlate more strongly with nuclear RNA-seq than with each other (Fig. S1A), indicating a more complex relationship between transcription initiation and nucleosome depletion. Discrete partitioning of genes into quintiles based on gene expression values derived from RNA-seq signal (1^st^ = lowest expression, 5^th^ = highest) further confirms that the relationships between nucleosome depletion, transcription initiation, and gene expression are imperfectly correlated. For instance, the highest gene expression quintile is characterized by reduced ATAC-seq enrichment as compared to the second highest quintile, despite it having the highest enrichment in Start-seq signal (Fig. S1B). These data demonstrate that open chromatin and transcription initiation do not directly track with each other.

We hypothesized that discrepancies between ATAC-seq, Start-seq, and nuclear RNA-seq could be due to the influence of promoter proximal polymerase pausing on the relationship between nucleosome depletion and transcription initiation, as has been shown previously [32]. Specifically, we sought to test whether polymerase pausing might increase the extent of nucleosome depletion within a NDR, as inferred from MNase-seq data in S2 cells [32]. To evaluate the relationship between differential pausing and chromatin accessibility, we derived ‘pausing index’ (PI) values for each gene by determining the ratio of TSS Start-seq signal vs. gene body nuclear RNA-seq signal. Whereas Start-seq and nuclear RNA-seq levels are correlated (Fig. S1A), a scatterplot of those values for each promoter identifies significant variability from the regression line, indicating a wide range of pausing propensities (Fig. 2A). Moreover, PI can predictably stratify classes of genes that are expected to be more (or less) paused on average, based on previous studies [33]. For example, many housekeeping genes exhibit very low PI values, consistent with their ubiquitous and temporally consistent expression (example in Fig. S2A). In contrast, immune response and transcription factor genes, which in many cases are subject to rapid temporal and signal-responsive regulation that is achieved by pausing, display high PI values (Fig. 2B, example in Fig. S2B). Furthermore, enrichment of the “Pause Button” *cis-* regulatory motif, which is characteristic of many paused promoters [34], is positively correlated with PI quartile (Fig. S2C). We conclude that the PI metric (as calculated here) is a biologically relevant measure of gene-specific pausing propensity.

**Figure 2:**
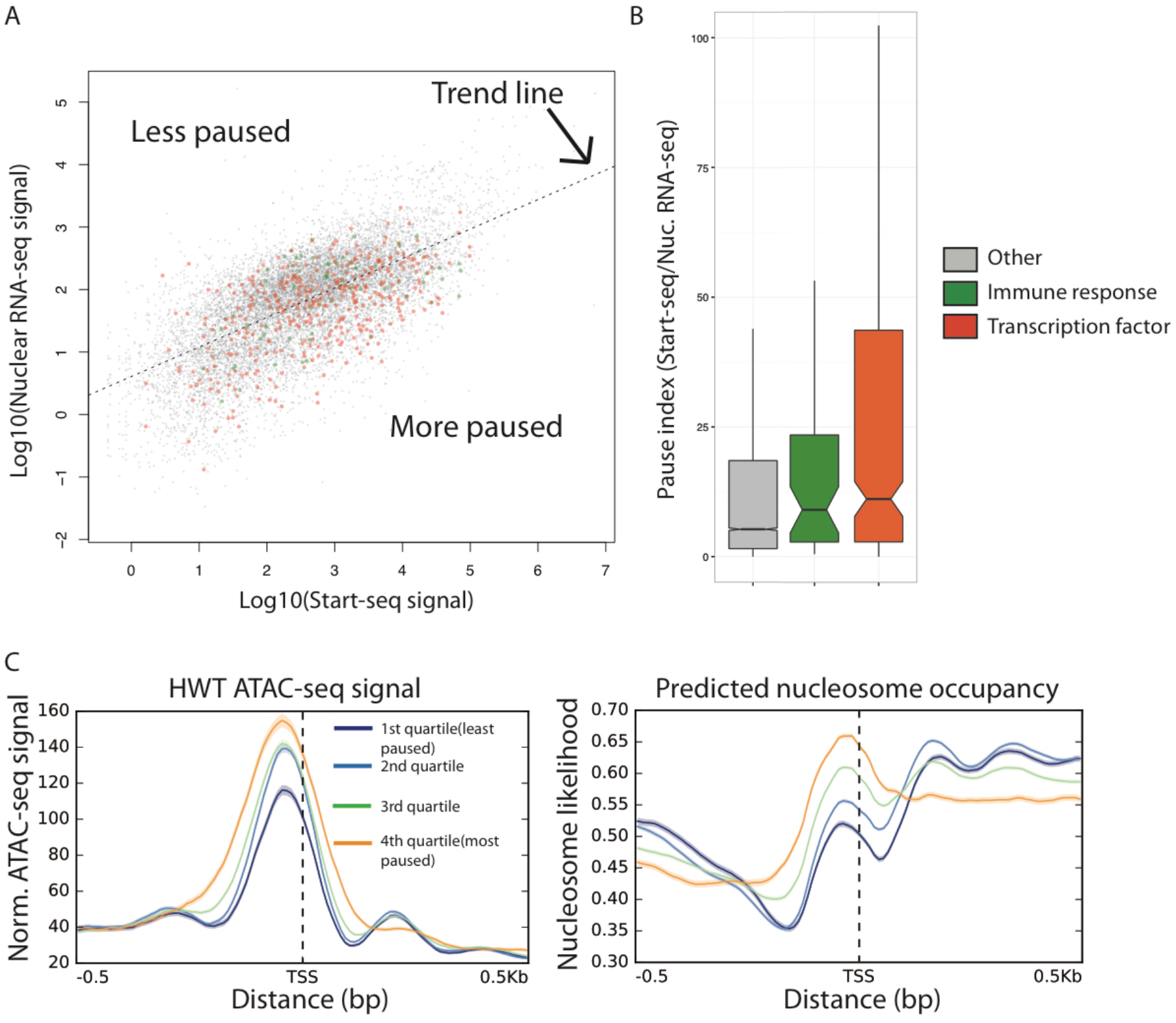
Relationship between polymerase pausing and chromatin accessibility. A) Scatterplot of Start-seq (x-axis) vs. nuclear RNA-seq (y-axis) signal. B) Pause index (Start-seq/RNA-seq) for obsTSSs from different classes of genes, including immune response and transcription factor genes (more paused than average). C) ATAC-seq signal and predicted nucleosome occupancy at obsTSSs stratified by pausing index (PI). High-PI obsTSSs have higher chromatin accessibility at the minus-1 nucleosome free region, despite also having higher predicted nucleosome occupancy in that region.

To test the relationship between pausing and chromatin accessibility, we partitioned genes into quartiles based on their PI values, and quantified ATAC-seq chromatin accessibility and predicted nucleosome occupancy in a window surrounding TSSs in those quartiles. We found that genes in the most highly paused quartile have the highest ATAC-seq signal at the minus-1 nucleosome position, despite also having the highest predicted nucleosome occupancy (Fig. 2C), which is consistent with previous observations [32]. Notably, the less-paused genes have a well-phased plus-1 nucleosome, further indicating that PI predicts the expected variability in nucleosome phasing based on pausing [32,35]. Using a direct assay for open chromatin in third instar larval nuclei, we conclude that polymerase pausing positively correlates with chromatin accessibility at the minus-1 nucleosome, in spite of underlying sequence information. These findings support the idea [32] that pausing may play an active role in maintaining NDRs.

### Start-seq signal clusters into spatially restricted groups of peaks at annotated TSSs

As observed in S2 cells [19], Start-seq signal often manifests as single-nucleotide peaks that are grouped in a spatially restricted region, such that TSSs can be described as “clusters” of initiation events at a handful of nucleotides near the 5’ end of a gene. To illustrate these clusters, we grouped individual +1 Start-seq nucleotides within 5nt of each other into likely TSS clusters, based on the fact that >50% of strong Start-seq peaks are within 5nt of the nearest neighbor peak (Fig. 3A), and assigned the clusters to annotated gene promoters. This procedure yielded 21,830 TSS clusters that matched stringent statistical criteria, 18,070 of which mapped to promoters annotated previously in the dm5.57 update of the *D. melanogaster* genome build. We termed these clusters <u>obs</u>erved promoter TSSs (obsTSSs). The remaining 3,123 high-confidence TSSs that failed to map to an annotated promoter region were considered <u>n</u>ovel <u>u</u>nannotated TSSs (nuTSSs).

**Figure 3:**
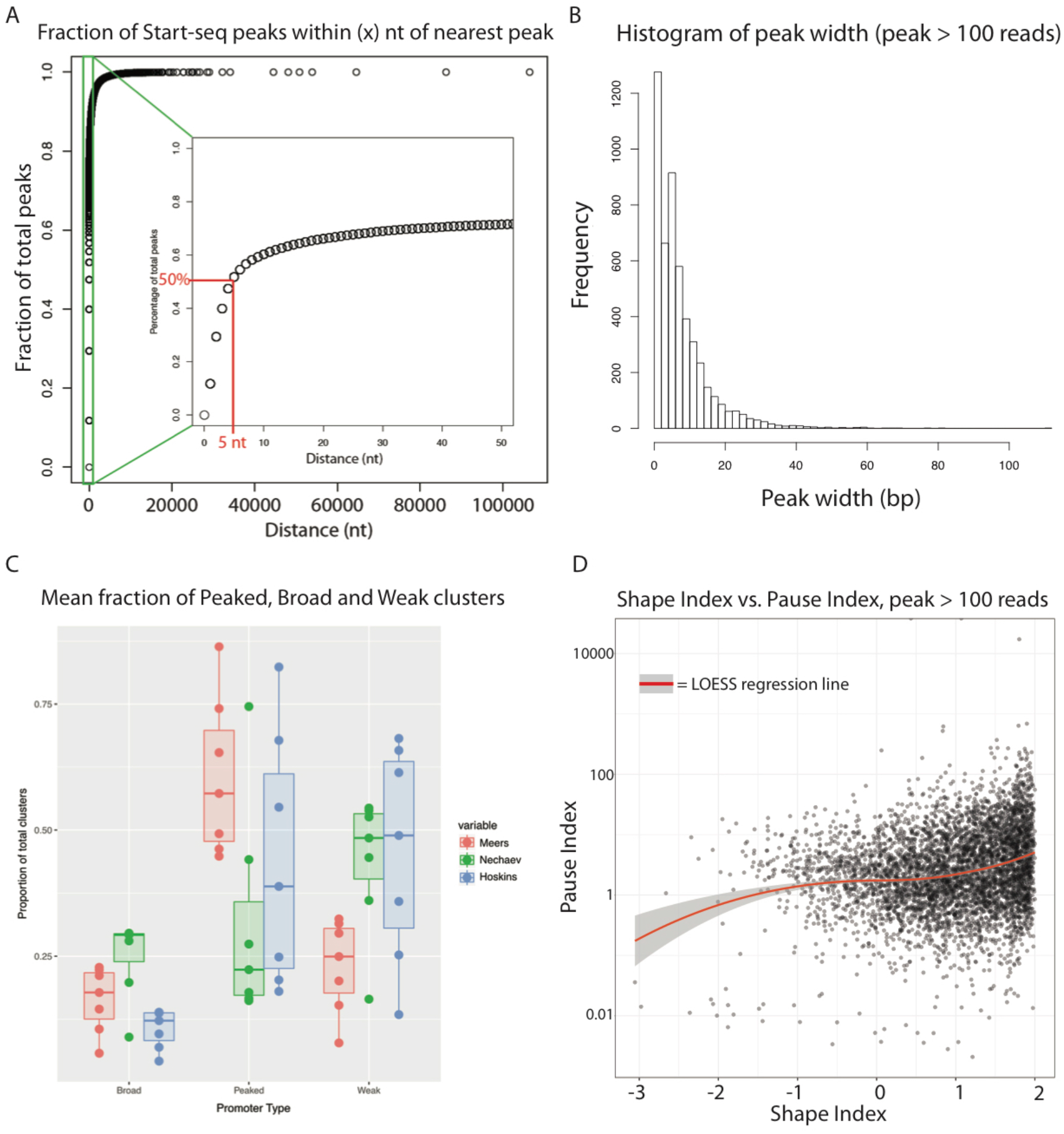
Characterization and comparison of TSS clusters. A) Cumulative percentage plot illustrating the fraction of Start-seq single nucleotide peaks within a given genomic distance of its nearest neighbor peak. Inset shows green region with expanded x-axis to illustrate that 50% of peaks are within 5 or fewer nt of the next nearest peak. B) Histogram showing the distribution of peak cluster widths for obsTSSs identified in Start-seq data. Notably, 50% of peaks are narrower than 6nt, in contrast with previously reported broader distributions of peak widths (Ni *et al.* 2010, Hoskins *et al.* 2011). C) Boxplot describing the proportion of broad (left), peaked (middle), and weak (right) peak clusters across a range of clustering thresholds from our data (Red), Nechaev et al. Start-seq data [19] (Green), and Hoskins et al. CAGE data [13] (Blue). For additional details, see Fig. S3D] Scatterplot comparing shape index (SI] with pausing index (PI).

Previous studies of *Drosophila* embryos and embryonic cell lines have revealed the presence of different TSS “shapes,” as defined by the breadth of the distribution of initiation signals within a given TSS [13,19,36]. Studies in mammalian cells have shown that TSS shape characteristics, particularly “sharp” and “broad” classifications, correlate with different sequence motifs enriched at the associated promoters, and different transcriptional outcomes from the corresponding genes [12,37]. To measure TSS shape in wandering 3^rd^ instar larvae, we measured cluster width and the fraction of total Start-seq signal in the cluster contained in the 5nt surrounding the highest peak in the cluster (Fig. 3B). Strikingly, maximum cluster width was substantially less than those values obtained by both Hoskins *et al.* [13] and Ni *et al.* [36], and more in line with S2 cell data from Nechaev *et al.* [19] who found that initiation was more highly focused. Specifically, among TSSs that were broader than a single nucleotide, ~47% of them were 6nt or fewer in width, ~75% were 10nt or fewer, and ~94% were narrower than 20nt. When we categorized TSSs as being “peaked” (<12nt in width), “broad” (>12nt, with > 50% signal in highest peak) and “weak” (>12nt, < 50%), we found that 81.8% of TSSs broader than 1nt would be classified as peaked, 8.0% as broad, and 10.2% as weak. Again, these numbers are quite distinct from the 32.6% peaked, 18% broad, and 49.4% weak, as described by Ni *et al.* [36].

We noted that the other groups employed different library preparation methods, and also relied on smoothing-density estimates for signal quantitation in peak clusters, and therefore their results may not be directly comparable. To determine whether library preparation or clustering parameters might have contributed to differences in “peaked” promoter identification, we directly compared the two methods. We subsampled reads from our Start-seq dataset, a Start-seq dataset from S2 cells [19], and a CAGE dataset from *D. melanogaster* embryos [13], such that the pairwise comparisons between our dataset and each of the two other sets used the same read depth, the same peak identification strategy, and the same peak merging distance thresholds (i.e. the largest distance between two peaks that can be considered part of the same cluster). Interestingly, across several distance thresholds, both the Hoskins et al. CAGE peaks and the Nechaev et al. [19] Start-seq peaks consistently under-represented “peaked” clusters and over-represented “weak” clusters relative to our peaks (Fig. 3C and Figs. S3A, S3B). This result argues against library preparation or cluster thresholding as confounding factors in comparing the sharpness of peaks, though it does not definitively rule out other technical differences. Nevertheless, given the significant depth to which we sequenced our libraries (collectively ~100M mapped reads), and the expectation that all peak shape modalities should be represented at that depth, we conclude that *D. melanogaster* TSSs are indeed largely “sharp” and focused, at least at the third instar larval stage. These results contrast with previous findings that broad TSSs are well represented in the fruit fly transcriptome but are consistent with the absence of CpG island promoters in *Drosophila,* a feature that is characteristic of less-focused TSSs in mammals [12].

### Cis-regulatory sequence motifs are enriched in patterns around obsTSSs that correlate with peak shape and polymerase pausing

Given our confirmation in larvae that *D. melanogaster* TSSs do not conform to the typical sharp/broad duality seen in mammals, we reasoned that TSS peak shape might correlate differently with sequence motifs and gene expression outcomes in flies. Therefore, we sought to generate a numeric shape metric that would allow us to elucidate relationships between TSS shape and other aspects of gene expression. To do so, we adapted the approach taken by Hoskins *et al.* [13] to assign a Shape Index (SI) value to each cluster (see Supplementary Methods), where higher SI values represent “sharper” peaks (i.e. the majority of TSS signal occurring within a few nucleotides), and lower SI values “broader” peaks (i.e. signal was spread more evenly across a wider locus). Because we found TSSs to be universally sharper than previously reported (and therefore more likely to have high SI values), most promoters had an SI value between -1 and +2 (Fig. 3D). Interestingly, shape index was mildly, but positively, correlated with pause index (Fig. 3D). This finding is consistent with the idea that high SI promoters are greatly enriched for the Pause button (PB) motif [13], and argues that SI remains a useful metric, despite the narrower overall distribution found in our study.

To determine whether peak shape or pausing index is associated with particular sequence motifs, we searched the regions flanking obsTSSs for a suite of motifs that were previously shown to be enriched at *Drosophila* promoters [38], and then clustered them based on motif enrichment (Fig. 4A). Unsupervised clustering analysis partitioned obsTSSs into three bins: a cluster characterized by the enrichment of GAGA, initiator element (INR), downstream promoter element (DPE), and PB (obsTSS Cluster 1); a cluster with reduced frequency of the aforementioned motifs and a strong enrichment of TATA (obsTSS Cluster 2); and a third cluster that was enriched for elements such as DRE and E-box and devoid of the other aforementioned motifs (obsTSS Cluster 3). obsTSS Cluster 3 had both the lowest PI and lowest SI among the three clusters (Fig. 4B, green), and is very similar to a previously identified class of low SI promoters that lack the PB motif [13]. obsTSS Cluster 1 has the highest PI (Fig. 4B), and corresponds to a class of high SI TSSs [13] enriched for many of the same motifs (GAGA, INR, PB). Similar to our observations of the effect of pausing on ATAC-seq signal relative to predicted nucleosome occupancy in Fig. 2C (highest PI quartile), the promoters in high-PI obsTSS Cluster 1 exhibited equally robust ATAC-seq signal to obsTSS Cluster 3 despite a higher predicted nucleosome occupancy (Fig. 4C). Interestingly, whereas both higher PI and SI distinguish obsTSS Clusters 1 and 2 from Cluster 3, #1 and #2 share similar SI values, further separating #3 as a functionally distinct class of broad TSSs with unique sequence elements.

**Figure 4:**
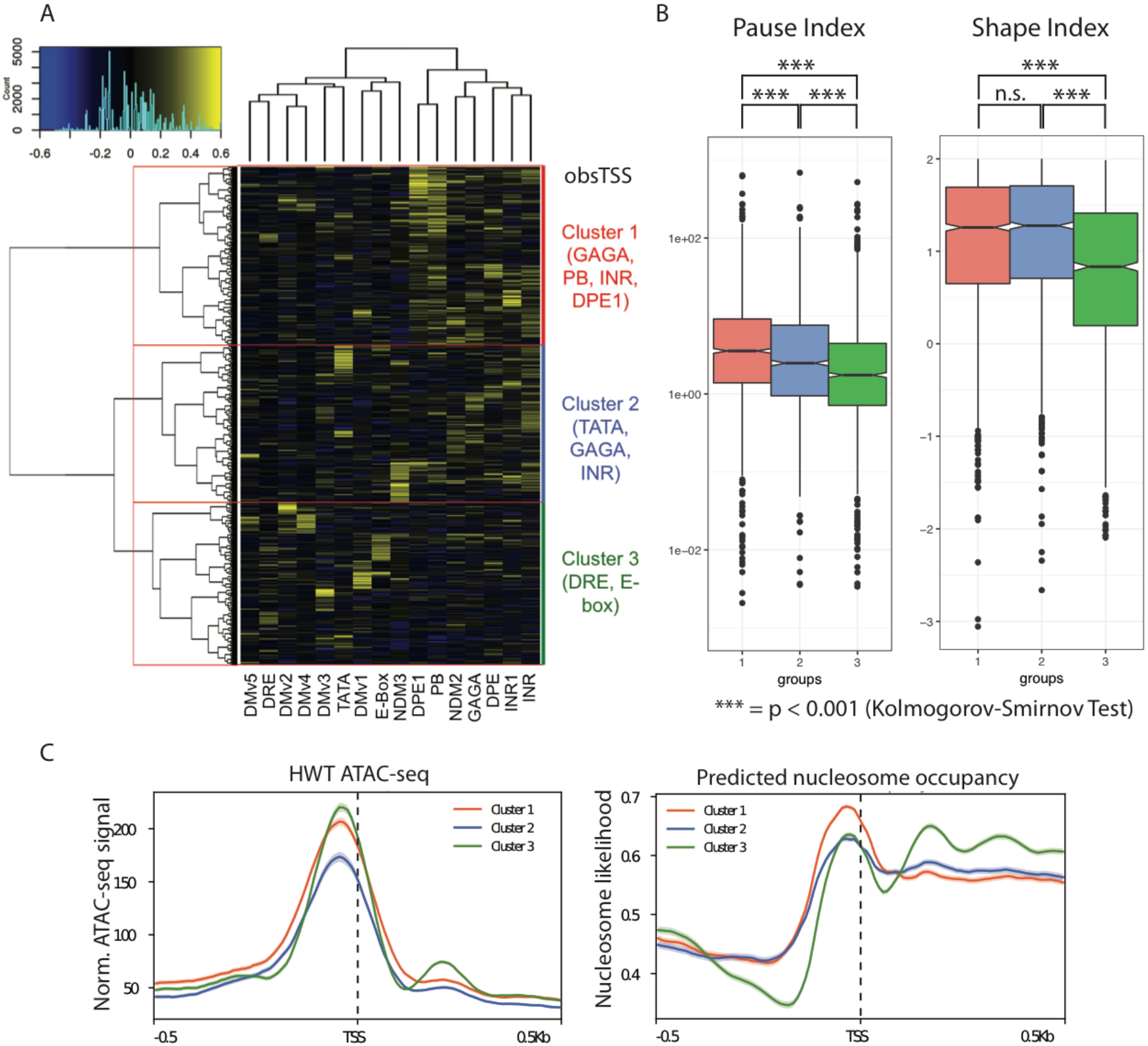
Analysis of TSS shape in *D. melanogaster* 3^rd^ instar larvae. Heatmap describing enrichment of 16 motifs associated with *D. melanogaster* promoters (columns] in each TSS (rows] with more than 100 start-seq reads mapping to its dominant peak. Row clustering dendrogram partitions genes into three groups (outlined in red, described at right). B) Boxplots describing distributions of Pause Index (left) and Shape Index (right) values in TSSs belonging to Cluster 1 (red), Cluster 2 (Blue), or Clutser 3 (Green). P-values generated by Kolmogorov-Smirnov Test. C) ATAC-seq signal (left) and predicted nucleosome occupancy (right) in a 1 kb window around obsTSSs belonging to Cluster 1, 2, or 3.

Interestingly, the motifs enriched in obsTSS Cluster 1 (and to a lesser extent Cluster 2) correlate positively with each other (Fig. S4A, dashed lines), indicating that multiple motifs tend to cooccur near the same TSS, whereas the motifs enriched in Cluster 3 are uncorrelated and tend to be mutually exclusive (Fig. S4A). This finding suggests that sharp, highly paused TSSs are more sequence-dependent than their broader, less paused counterparts. Consistent with this hypothesis, in addition to being enriched for several known motifs both upstream and downstream of its member TSSs, obsTSS Cluster 1 had the highest information content in its consensus sequence directly surrounding the TSSs, implying a role for sequence in defining the characteristics of sharp TSSs (Fig S4B). Further, 67% of promoters that can be considered “sharp” in embryos remain so in larvae, whereas only 27% of peaks considered “broad” remain that way in larvae, suggesting that intrinsic cis-regulatory information contributes to sharp, but not to broad promoters. By using stages of *Drosophila* development that have not been previously analyzed, we show that transcription factor and other cis-regulatory motifs are reliably correlated with TSS shape and polymerase pausing. Taken together with previous work [12,13,36], our data provide strong support for functional connections between sequence motifs, promoter-proximal pausing and TSS shape across multiple developmental points in *Drosophila.*

### Divergent promoters in *D. melanogaster* larvae

“Divergent” promoters, regions from which transcription proceeds from a coupled set of core promoter elements oriented in opposite directions, have been widely reported in mammalian systems [14,15,17]. Antisense transcription from divergent TSSs is thought to have regulatory consequences for expression of the sense-oriented protein-coding gene [17]. However, there is very little evidence of the same phenomenon in *D. melanogaster* [19], though to date it has not been analyzed *in vivo* in an organismal context. Using our high-depth 3^rd^ instar larval Start-seq dataset, we searched for divergent transcription units. For a given TSS, we mapped the fraction of Start-seq signal accumulating in sense and antisense directions relative to each site in question. We found that both obsTSSs and nuTSSs exhibited highly sense-oriented signal (Fig. S5A). We also quantified sense-oriented reads as a proportion of the total reads mapping in a 200nt window on either side of each TSS. Although obsTSSs were statistically more enriched for sense-oriented reads than were nuTSSs, the mean was greater than 90% sense-oriented for both cohorts, indicating a high degree of unidirectionality at all TSSs in *D. melanogaster* (Fig. S5B).

To identify divergent TSSs, we aligned all high-confidence TSSs whose nearest neighboring TSS was oriented in the opposite direction, then ordered them based on genomic distance between each pair, and plotted ATAC-seq signal in order to identify single NDRs housing the two TSSs. A threshold distance of roughly 200nt between the TSS pair yielded 537 pairs of TSSs for which a single continuous ATAC-seq NDR overlapped both TSSs (Fig. 5A). These 537 pairs contained 1,023 distinct TSSs, or ~4.8% of the all the high-confidence TSSs queried (Fig. S5C, example in Fig. 5B), as compared with greater than 75% of active promoters observed with divergent transcription in mammalian systems (Scruggs et al. 2015). Despite the dearth of paired TSSs genome-wide, of those that were paired, 444 (43.4%) were obsTSSs paired with another obsTSS from separate, divergent coding genes, which we refer to as bidirectional promoters (Fig. S5C). This is in contrast with the human transcriptome, in which only a small proportion of divergent transcription is represented by bidirectional promoters [39].

**Figure 5:**
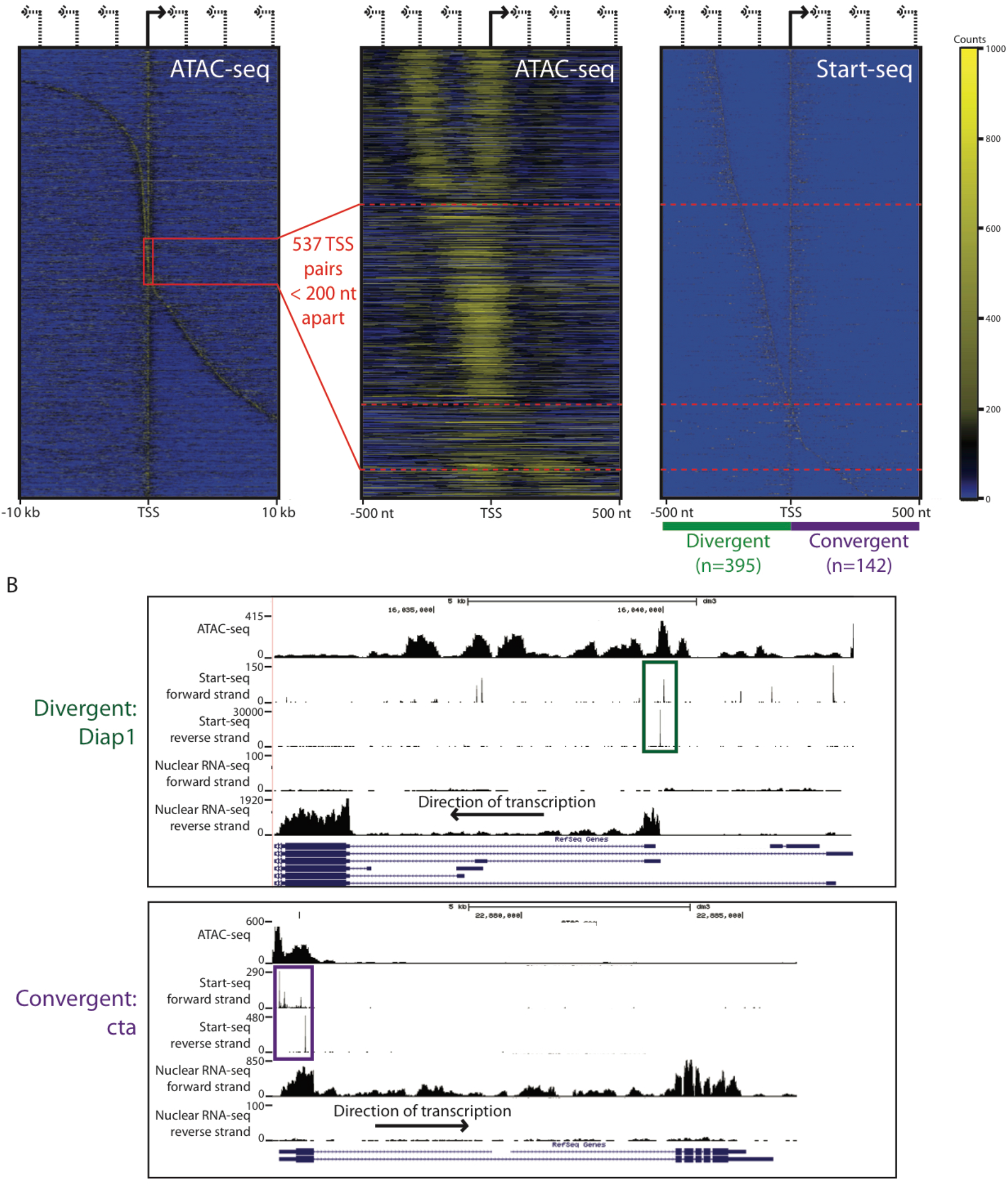
Divergent transcription in *D. melanogaster*. A) Panel 1 (left): Heatmap of ATAC-seq signal mapping in a 20 kb window around TSSs whose nearest neighbor TSS is oriented in the opposite direction. TSSs are ordered by distance between TSS pair. Panel 2 (center): Heatmap of ATAC-seq signal mapping in a 1 kb window around TSS pairs separated by less than 300nt. TSS pairs separated by less than 200nt (highlighted by red box in panel 1) are partitioned into divergent or convergent TSS pairs by red dashed lines. Panel 3 (right): Heatmap of Start-seq signal for TSSs exhibited in panel 2. B) Representative browser window examples of divergent (top) and convergent (bottom) obsTSS pairs.

Divergent transcription might aid in recruiting transcription initiation machinery and maintaining a robust NDR at active promoters. To determine whether nucleosome depletion is increased at sites of divergent initiation in *Drosophila,* we quantified ATAC-seq reads in a 200nt window around obsTSSs participating in bidirectional, divergent, or non-divergent initiation, and compared it to Start-seq levels (Fig. S5D). We found that bidirectional obsTSSs had significantly more ATAC-seq signal than divergent or non-divergent obsTSSs, despite the expectation that ATAC-seq would correlate with the lower level of Start-seq signal at bidirectional obsTSSs (Fig. S5D). However, it is known that bidirectional promoters are often separated by the BEAF32 insulator in *Drosophila* [40], and indeed bidirectional promoters were enriched for BEAF32 ChIP-seq signal relative to divergent and non-divergent TSSs (Fig. S5E). Therefore, we could not rule out the possibility that increased ATAC-seq signal at bidirectional promoters may be due to displacement of nucleosomes by BEAF32. Importantly, divergent and non-divergent obsTSSs exhibited similar levels of BEAF32 ChIP-seq signal (Fig. S5E), and negligible differences in ATAC-seq signal (Fig. S5D), indicating that divergent transcription is generally insufficient to enforce a more robust NDR than would be expected by initiation activity in *D. melanogaster.* This observation is consistent with the finding that RNA pol II and H3K4me3 ChIP-seq accumulation is similar between directional and divergent TSSs in S2 cells [41].

Strikingly, 142 of the TSS pairs we identified were not divergent, but rather were oriented towards each other (Fig. 5A, example in Fig. 5B). We termed these “convergent” pairs, and they included 86 obsTSSs converged on by nuTSSs, and 16 pairs of obsTSSs that converged and productively elongated in both directions. In general, the distance between convergent pairs of TSSs was much larger than that of divergent pairs, indicating selection against convergent transcription in close genomic proximity (Fig. S5F). These results are consistent with the characteristics of convergent initiation pairs detected in mammalian cell culture [42], and confirm the presence of convergent transcription in an *in vivo* context. However, similarly to other studies, we cannot rule out the possibility that convergent transcripts originate from distinct cell populations. We conclude that, within a broader regime of unidirectionality, several *D. melanogaster* TSSs represent striking exceptions.

### Novel unannotated TSSs (nuTSSs) are widespread and can be partitioned into predicted functional categories based on local histone modifications

Owing to the depth of our Start-seq libraries (>100M mappable reads combined), we were able to identify bona-fide Start-seq signal at thousands of locations across the genome that did not correspond to an annotated TSS. To systematically analyze these locations, we applied several metrics to peaks that did not fall in a TSS cluster that matched to an existing obsTSS (nuTSSs, as defined previously). We identified a total of 11,916 distinct nuTSSs, including 3,123 that met a relatively stringent false-discovery-rate (FDR) threshold of 9 reads within every biological replicate. In general, nuTSSs exhibit NDRs comparable in shape to those found at obsTSSs, despite residing at loci with a higher intrinsic likelihood of nucleosome occupancy (Fig. S6A). nuTSSs are spread throughout the genome, although they predominantly cluster within, or proximal to, annotated genes (Fig. S6B).

A handful of well characterized histone post-translational modifications (PTMs) co-localize with bona-fide TSSs, and therefore we surmised that enrichment of particular histone PTMs at nuTSSs might provide an indication of nuTSS functions. Therefore, we measured the enrichment of a battery of histone PTMs at nuTSSs, and conducted unsupervised hierarchical clustering. Out of several modEncode ChIP-seq tracks (www.modencode.org) taken at a matched stage of development, we found that the most informative set of PTMs included H3K4me1, H3K4me3, H3K27ac, and H3K36me3. Hierarchical clustering based on these four marks resulted in seven categories (Fig. 6A). We characterized them as follows: a “Featureless” group (nuTSS Cluster 5) lacking significant enrichment in any of the four marks, “TSS-like” groups (nuTSS Clusters 1, 2, and 6) with enrichment for H3K4me3 characteristic of annotated coding gene start sites [43], two “Coding” cohorts (nuTSS Clusters 3 and 4) characterized by varying levels of enrichment for H3K36me3 as is expected in gene bodies, and an “Enhancer-like” group (nuTSS Cluster 7) with enrichment of H3K4me1 and H3K27ac marks characteristic of enhancer regions [44–46]. We further validated these functional classifications by observing that the “active” cohorts (TSS-like and Enhancer-like) were generally accompanied by strong ATAC-seq signal comparable to that of obsTSSs, whereas the other cohorts were depleted of ATAC-seq signal (Fig. 6B). Strikingly, the majority of nuTSSs clustered into “Featureless” (1296/3123, 41.5%) or “Enhancer-like” (715/3123, 22.9%) cohorts that lacked H3K4me3, indicating the prevalence of transcription initiation events that may serve functions other than transcription of as-yet unannotated genes. Together, these findings strongly suggest that nuTSSs localize within functionally relevant chromatin contexts across the genome.

**Figure 6:**
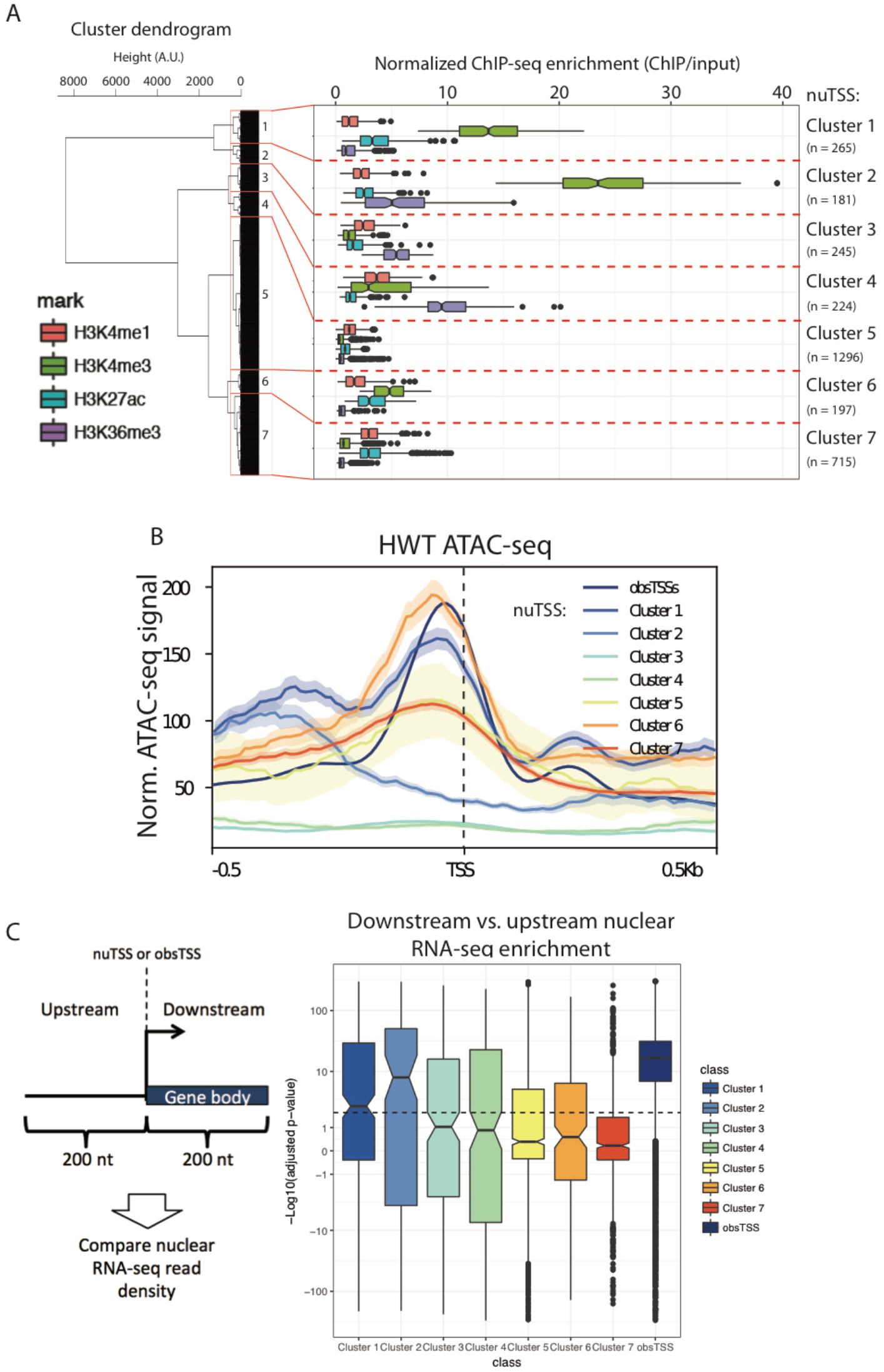
Novel unannotated TSSs (nuTSS) cluster into functional categories. A) Clusters of nuTSSs based on enrichment of H3K4me1 (red), H3K4me3 (green), H3K27ac (teal), and H3K36me3 (purple). The number of nuTSSs in each cluster is indicated at the right of the plot. B) ATAC-seq signal at obsTSSs (dark blue) and nuTSSs from different histone PTM-based clusters. C) Nuclear RNA-seq reads were mapped to regions 200nt upstream and downstream of each TSS, and enrichment of downstream vs. upstream reads was analyzed as a proxy for elongation. At right: boxplot of -Log10-transformed adjusted p-values for downstream signal enrichment over upstream within each nuTSS cluster (downstream-enriched nuTSSs above zero, upstream-enriched nuTSSs below zero).

### Most nuTSSs do not produce stable transcripts

Although high-throughput sequencing methods have enabled extensive and detailed annotation of global transcriptomes, studies that employ ever higher depth and sensitivity are continuing to uncover previously undiscovered RNAs. The high depth of our Start-seq libraries may have identified initiation sites for unannotated genes that undergo productive elongation and produce mature, stable transcripts, particularly for nuTSSs associated with PTM-based clusters enriched for H3K4me3. To determine whether this was the case, we mapped nuclear RNA-seq reads in 200nt windows upstream and downstream of all nuTSSs, and measured the balance of signal on either side, reasoning that productive elongation would be identified by an overrepresentation of downstream reads. We confirmed this hypothesis by performing the same test on obsTSS, and found that they are universally enriched for RNA-seq reads in the downstream region over the upstream (Fig 6C, Fig. S6C). In contrast, nearly all nuTSSs showed no significant enrichment of downstream signal (Fig. S6C). This trend held for most of the histone PTM-defined nuTSS clusters, where all but Clusters 1 and 2 had a mean adjusted p-value of downstream signal enrichment that was below the minimum threshold for significance. (Fig. 6C, dashed line). This observation suggests that nuTSSs generally are not converted into mature, stable transcripts.

Because Clusters 1 and 2 were most biased towards downstream vs. upstream read density, and were both highly enriched for H3K4me3, we investigated them more closely to determine whether their constituents represented unannotated “canonical” TSSs that resulted in elongating mRNAs. nuTSS Cluster 1 in particular uncovered distinct cases wherein the updated dm6 *D. melanogaster* genome build annotated additional first exons that were overlapped by Cluster 1 nuTSSs derived from an earlier build (example in Fig. S6D). When we intersected our nuTSSs with 5’UTR regions converted from the most recent genome build, 37 nuTSSs in Cluster 1 overlapped, suggesting that several Cluster 1 nuTSSs in fact correspond to mRNA initiation sites. Overall, we find that nuTSSs do not elongate into stable transcripts, with few exceptions corresponding to newly identified coding gene start sites.

### nuTSSs enriched for enhancer-associated chromatin marks overlap with functionally validated, tissue-specific enhancers

Recent studies in mammalian model systems have reported the presence of short-lived transcripts (eRNAs) originating from developmentally-regulated enhancers [14,15,17]. These findings are evocative of regulatory non-coding RNAs (ncRNA) at enhancer regions in *Drosophila* [47]. As mentioned above, classification of nuTSSs by histone PTMs showed that nuTSS Cluster 7 is distinguished by local enrichment of H3K4me1 and H3K27ac, both of which are hallmarks of active enhancers (Fig. 6A). We therefore termed nuTSSs belonging to Cluster 7 “Enhancer-like nuTSSs” (E-nuTSSs). As with previously reported eRNAs, E-nuTSSs are associated with robust NDRs (Fig. 6B), suggesting assembly of the PIC similar to “canonical” promoters. E-nuTSSs predominantly appear within introns (~74%), whereas only ~12% occur in coding sequences (Fig. 7A), which is suggestive of low sequence conservation and perhaps more recent cis-element evolution. In agreement with this interpretation, the information content of the E-nuTSS consensus sequence is low, and comparable to that of lowest information cohort of obsTSSs that we analyzed (Fig. 7B).

**Figure 7:**
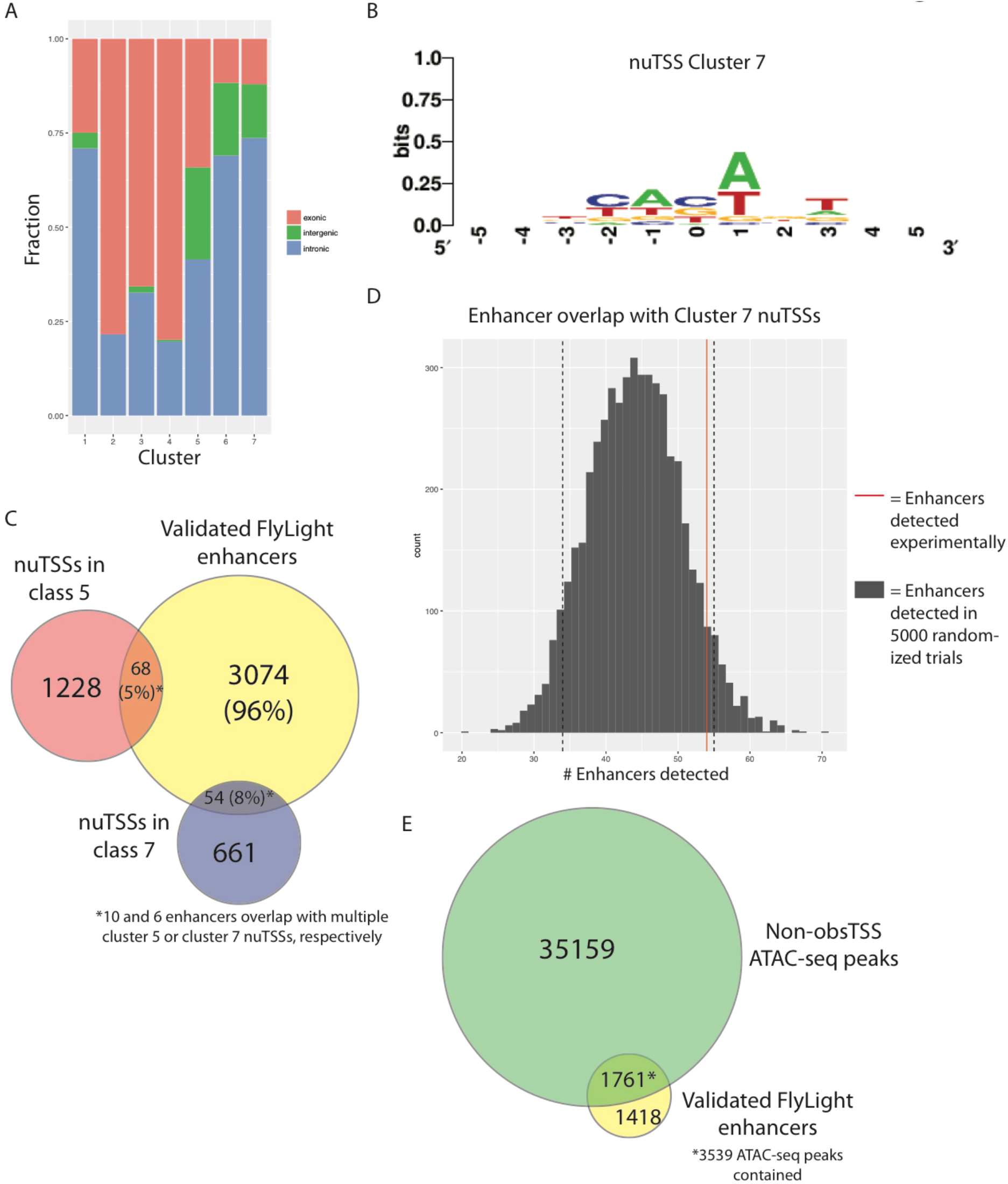
Analysis of E-nuTSS distribution and overlap with known enhancers. A) Genomic distribution of nuTSSs in different histone PTM-based clusters. “Enhancer-like” (nuTSS cluster 7) start sites occur disproportionately in intergenic and intronic regions. B) Consensus motif at Cluster 7 nuTSSs. C) Overlap between Flylight enhancers validated for expression in larval CNS or imaginal discs (yellow, 3179) and nuTSSs from Cluster 5 (red, 1296) or cluster 7 (blue, 715). D) Histogram describing number of nuTSSs overlapping validated FlyLight enhancers in 5000 randomly shuffled trials (grey) vs. the number of Cluster 7 nuTSSs overlapping experimentally (red line). E) Overlap between Flylight enhancers validated for expression in larval CNS or imaginal discs (yellow, 3179) and ATAC-seq peaks (Green, 38698).

If E-nuTSSs represent bona-fide eRNAs, they would expand the ensemble of known regulatory RNAs in *D. melanogaster,* a model system wherein eRNAs have not been characterized systematically. To determine whether E-nuTSSs overlap with functional enhancer regions, we curated *D. melanogaster* 3^rd^ instar larval enhancers from the FlyLight collection [48,49]. These enhancers have been functionally validated via GAL4-UAS based reporters, and we chose 3,179 of the 7,113 total enhancers in the collection on the basis that they were shown to promote expression of a fluorescent reporter protein in either larval imaginal discs or in the larval CNS. Among the 3,123 high-confidence nuTSSs interrogated, we found that 135 unique high-confidence nuTSSs overlapped directly with one of the 3,179 validated enhancer regions, and that 116 enhancers contained at least one nuTSS. These numbers represented ~4.4% and ~3.6% of the populations queried, respectively (Fig. 7C). Importantly, 90% (122/135) of the nuTSSs that overlap with an enhancer belonged to either Cluster 5 (68, 5% of all Cluster 5 nuTSSs) or Cluster 7 (54, 7.6%). When Cluster 7 regions were randomized throughout the genome prior to measuring overlap with enhancer regions *in silico,* fewer of the resultant shuffled nuTSSs overlapped with enhancers than did Cluster 7 nuTSSs in 94% percent of 5000 random trials (Fig. 7D). Thus histone PTM-derived clustering analysis is useful for identifying potential eRNAs.

The low fraction of enhancers that overlapped with high-confidence E-nuTSSs indicates that TSS profiling alone is unlikely to be useful as a tool for predicting enhancers as compared with other methods. For instance, open chromatin data have been used to detect tissue-specific enhancers in *D. melanogaster* [50]. We therefore used a set of 38,696 ATAC-seq peaks that do not contain an obsTSS to measure overlap with enhancers, and found that 3,537 peaks (~9.1%) overlapped with 1,761 known enhancers (i.e. 55% of the total enhancer set; Fig. 7E). Notably, ATAC-seq peaks that overlap with enhancers were more likely to contain Start-seq signal than either randomized regions, or ATAC-seq peaks that do not overlap enhancers (Fig. 8A). Thus, our data reveal a clear propensity for E-nuTSSs (eRNAs) to occur within enhancer NDRs. We conclude that *D. melanogaster* enhancers are generally characterized by nucleosome depletion, and that these NDRs are more likely to be transcribed than other regions of comparable chromatin accessibility. However, at our present sequencing depth, and in a heterogeneous cell population, we find only a small fraction of these enhancer NDRs are associated with highly expressed eRNAs.

**Figure 8:**
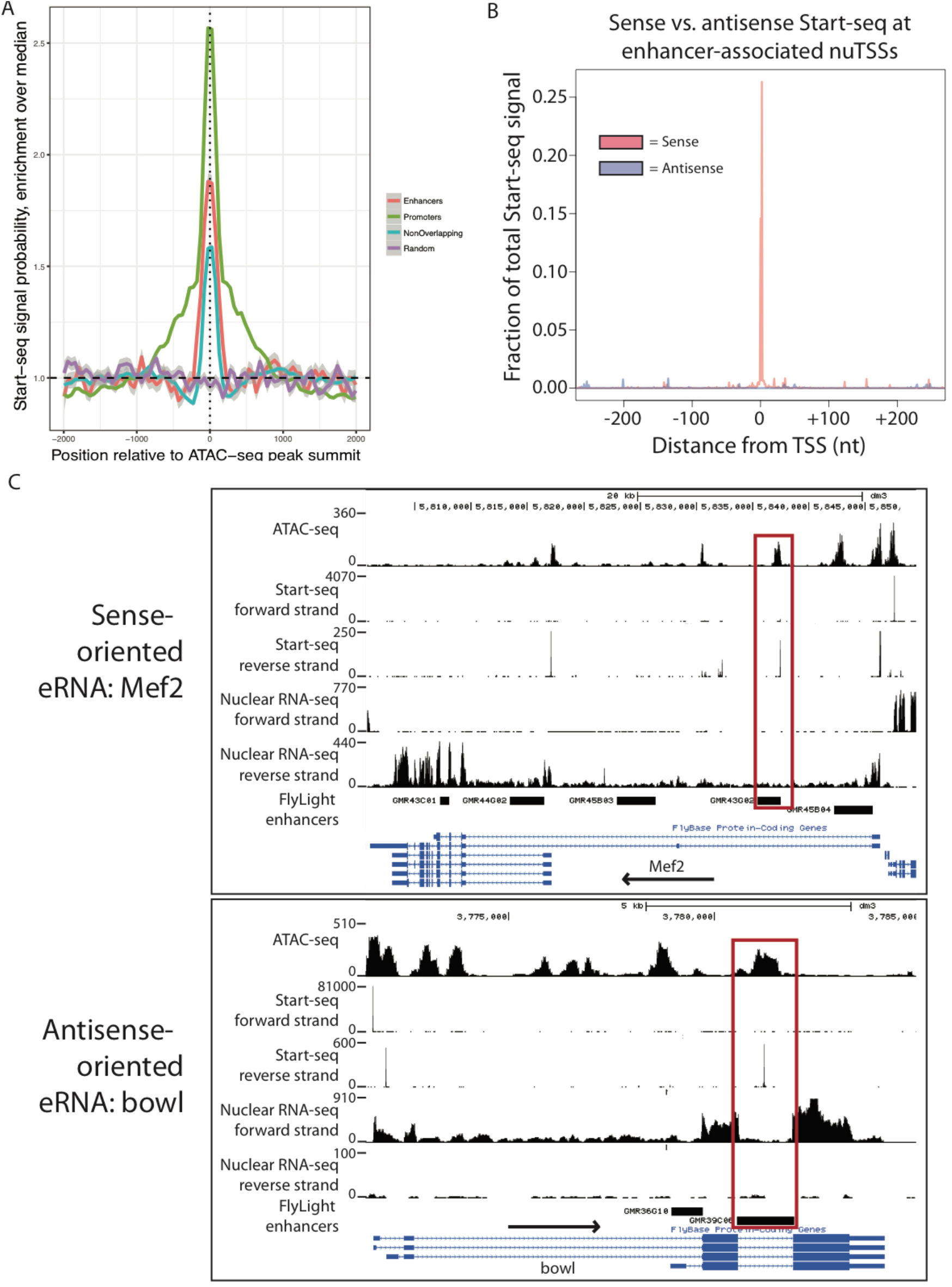
Detection of enhancer RNAs from Start-seq data. A) Metaplot describing position-specific enrichment over full window median probability of Start-seq signal in a 4 kb window around summits of ATAC-seq peaks that overlap with gene TSSs (Promoters, green), overlapping FlyLight enhancers (Enhancers, red), or overlapping neither feature (Non-overlapping, blue), along with randomly assorted genomic regions (Random, purple). B) Metaplot of sense (red) or antisense (blue) Start-seq signal mapping in a 500nt window around enhancer-associated nuTSSs. C) Representative browser window examples of enhancer-associated nuTSSs oriented in the sense (top) or antisense (bottom) direction relative to the gene containing the enhancer.

### nuTSSs associate with broadly expressing enhancers and represent unidirectional eRNAs

We hypothesized that the highly expressed eRNAs we were able to detect might be located within enhancers that are active in a large fraction of cells. Therefore, we examined expression data for the enhancer regions that contained nuTSSs compared to those that did not. Relative to the 3,063 enhancers that did not contain a high-confidence nuTSS, enhancers that contained a nuTSS were more likely to be enriched for expression in all five imaginal disc categories (leg, wing/haltere, eye, antennal, and genital) as well in the optic lobe, all of which represent large, broad-based cell populations (Fig. S7A). Similarly, nuTSS-containing enhancers were depleted for expression in various neural lineages, including brain-, subesophageal- and thorax-specific neurons, all of which are derived from small cell populations requiring highly specific regulatory elements (Fig. S7A). Furthermore, 26% (30/116) of nuTSS-associated enhancers express in 5 or more distinct tissues as compared with 8.3% (253/3063) of other enhancers. These data suggest that either eRNAs are primarily associated with broadly expressing enhancers, or that our stringent threshold (≥9 reads in each biological replicate) for nuTSS identification was too high to detect eRNAs expressing in a smaller number of cells. In the latter case, we reasoned that lowering the threshold for nuTSS detection would uncover overlap with additional enhancer regions. To distinguish between these possibilities, we intersected the remaining 8,793 nuTSSs that met a minimum statistical (FDR) threshold for detection, but not our more stringent threshold, with the base set of 3,179 enhancers. The low-stringency nuTSSs overlapped with 572/3179 total enhancers (~18% of the base set), including 456 of the previously unrecognized ones. Among the newly detected enhancers, only ~15.1% (69/456) of them expressed in five or more distinct tissues, indicating that eRNA expression levels are proportional to overall enhancer usage. These findings are consistent with the observed higher probability of Start-seq signal occurring within enhancer-overlapping ATAC-seq peaks (Fig. 8A). Thus, eRNAs may well be expressed at many more enhancers than those identified here, as our current detection threshold represents a lower limit.

Although eRNAs in mammalian cells are typically divergently transcribed [14,15,17,51], we find no evidence of divergent transcription from validated enhancers that overlap with a nuTSS, indicating that most eRNAs in *D. melanogaster* conform to the unidirectional characteristic of obsTSSs (Fig. 8B, Fig. S7B). Of the 135 high-confidence nuTSSs that overlap with validated enhancers, only nine of them participate in a divergent transcription pairing. Intriguingly, unidirectional eRNAs can be oriented in either the sense or antisense direction relative to their resident gene (75 sense vs. 51 antisense, examples in Fig. 8C). In summary, we conclude that eRNAs in *Drosophila* are unidirectional and correlated with enhancer strength.

## Discussion

Fidelity of transcription initiation is crucial for proper regulation of gene expression. There are several characteristics of transcription from annotated promoters that are thought to correlate with aspects of downstream gene expression, including promoter nucleosome depletion [11], interaction of initiation complexes with cis-regulatory motifs, start site “shape” [12], and promoter-proximal pausing [19,20,32]. To analyze transcription initiation in *D. melanogaster,* we performed matched ATAC-seq, Start-seq, and nuclear RNA-seq in larvae. For accurate developmental staging, we selected animals displaying the wandering behavior that is characteristic of the late 3^rd^ instar. We elucidated regulatory trends for initiation at annotated coding genes genome wide that are largely in agreement with previous studies, and also uncovered myriad sites of unannotated transcription.

### The role of polymerase pausing in NDR establishment

Previous studies have shown that promoter proximal polymerase pausing can have an effect upon local chromatin structure [32]. Specifically, highly paused genes are equally depleted of nucleosome signal at their promoters as their less paused counterparts, despite having a higher likelihood of promoter nucleosome occupancy based on sequence information alone [32]. Here we use a direct measure of chromatin accessibility (ATAC-seq) rather than nucleosome occupancy to show the same discrepancy at highly paused genes in third instar larvae (Fig. 2C). This discrepancy holds true even when TSSs aren’t explicitly grouped by pause likelihood. For instance, when we clustered TSSs by motif density, the cluster of TSSs with the highest median pause index exhibited the same average ATAC-seq signal as the cluster with the lowest PI, despite having a much higher predicted nucleosome occupancy (Fig. 4C). Based on these findings, we concur with a previous interpretation suggesting that pausing facilitates chromatin accessibility at promoters where nucleosome assembly is intrinsically favored [32]. We also offer this as a potential explanation for the unexpectedly low correlation between Start-seq and ATAC-seq signal at TSSs. To date, no systematic, direct analysis of chromatin accessibility paired with TSS signal from techniques such as CAGE or PEAT has been reported, and thus the hypothesis that the two measures should correlate well is based primarily upon mechanistic assumptions. Though we cannot rule out other biological contributions to the low correlation we report, the previously suggested Pol II pausing-mediated mechanism [32] that is directly supported by our data is the most parsimonious explanation.

### Relationship between peak shape and transcriptional outcomes

Previous TSS mapping studies in *Drosophila* [13,19,36] and other organisms [12] noted that the overall shape of a TSS domain had potential functional implications. We find that *D. melanogaster* larval TSS peaks are typically very sharp and focused. We also find that broad TSS tend to occur at highly expressed, lowly paused genes, whereas sharp peaks are highly paused and enriched for a host of cis-regulatory elements. Importantly, the average width of TSSs we detect from larvae is in contrast with TSSs from embryos, in which broader TSSs are more prevalent [13,36]. We initially attributed this difference to the library preparation techniques used in the various studies (Start-seq (this study); PEAT [36]; and CAGE [13]). However, the discrepancy between global peak widths is consistent even when controlling for read depth, library preparation, and peak detection and clustering strategies (Fig. 3C, Fig. S3). It is possible that the particular developmental stage at which we observed TSSs (third instar larval stage) vs. those of other studies (embryos [13,36]; and S2 cells [19]) could account for the difference, but more experiments will be required to evaluate this hypothesis. Nevertheless, despite the fact that the average width of our obsTSSs is considerably smaller than the minimum width detected using other methods [13], we were still able to stratify TSSs into functional categories based on the width and distribution of reads within a given domain or peak. Thus, even if *D. melanogaster* TSSs are not similarly “sharp” across all developmental stages and cell types, the functionality of differently shaped TSSs may be retained.

Interestingly, the broad TSSs that we infer to lack stably paused Pol II could be considered analogous to similarly broad mammalian housekeeping promoters that are enriched for CpG islands [37]. Given the absence of CpG island promoters in *Drosophila,* this finding suggests a convergent evolutionary force that promotes a broad modality of transcription initiation specifically at ubiquitously expressed genes. Although direct comparisons of peak width across sequencing platforms are difficult, most of the peaks that are considered “sharp” in embryos are also considered sharp in larvae, whereas the shape of broad peaks is less well conserved. Combined with the more consistent sequence context of sharp promoters (Fig. S4B), it is possible that sharp peaks were selected during evolution to behave as such because of the necessity of promoter proximal pausing-related regulation of their expression, whereas broad peaks require no such constraints, and are instead driven by their strong propensity for nucleosome depletion. Whether or not TSS peak shape is a functional characteristic upon which natural selection can act is speculative and will require further study.

### The role of promoter directionality in *D. melanogaster*

Previous studies have found that *D. melanogaster* promoters are highly unidirectional [19], despite voluminous data that argues for intrinsic bidirectionality of promoters in mammalian systems [14–17]. However, the unidirectional character of *Drosophila* TSSs has not been evaluated carefully in endogenous tissues. Here we find that similar to previous studies, TSSs lack antisense transcription initiation for the vast majority of promoters in wandering 3^rd^ instar larvae. However, we also uncover hundreds of new cases of divergent transcription, which we conservatively define as transcription initiating in two directions from the same contiguous nucleosome free region (NDR). Notably, more than half of the divergent TSS pairs we identified correspond to a bidirectional promoter pair in which annotated genes are transcribed in opposite directions from the same promoter region. There is evidence of this phenomenon in the human transcriptome [39,52], but it represents a small proportion of all divergent TSSs.

It has been suggested that divergent transcription may serve to strengthen the recruitment of transcription factors and other initiation machinery to the site of the sense-directed gene, thereby increasing its expression [53]. Though we detect much higher ATAC-seq signal at bidirectional obsTSSs than at unidirectional obsTSSs (Fig. S5D), it is unclear whether this is due to a synergistic effect of coordinated recruitment of transcriptional machinery, whether it reflects the fact that more cells are initiating from one TSS over the other, or whether the complex effects of insulator binding upon observed ATAC-seq signal are at play.

### Identification and characterization of enhancer RNAs

We showed that around 18% of validated larval enhancers from the Janelia FlyLight collection [48,49], also overlapped with a nuTSS peak identified in our Start-seq experiments. Although this number likely represents a lower bound, to our knowledge, no exhaustive post-hoc functional validation of a set of predicted enhancers has yet been undertaken. Hence, it is unclear whether our findings are comparable to other genome-wide approaches that have used enrichment of histone post-translational modifications (PTMs) [44,46], transcription factor binding sites [54], or regions of open chromatin or DNase hypersensitivity [50] to identify enhancers.

From our ATAC-seq data, we find that NDRs are more successful at detecting validated enhancers than are nuTSSs alone. From a practical perspective, using nucleotide-resolution TSSs to identify developmentally regulated enhancers may have the further benefit of improving the resolution of enhancer identification. Though enhancer length is variable, and the functional fraction of a given enhancer is undoubtedly longer than the spatially restricted region identified by a nuTSS, single-nucleotide resolution provides a helpful starting point for defining minimal sequence requirements for activation. It remains to be seen whether sequencing nuTSSs to higher depth could reliably identify spatially restricted enhancers. Future studies that carefully benchmark enhancer detection methods with experimentally validated enhancers will be instructive in this regard. Furthermore, it is possible that incorporating nuTSSs with other existing methods of enhancer detection, such as nucleosome depleted regions, may improve our ability to identify novel enhancers, particularly those that are active broadly in several tissues within a complex mixture of cells. Indeed, our annotation of nuTSSs with their enrichment for enhancer-associated histone PTMs already partially achieves this goal.

### Assaying eRNAs in *D. melanogaster*

From the perspective of *D. melanogaster* eRNA function, it is important to note that Start-seq signal was generally more likely to occur within enhancer-overlapping NDRs than it was at comparable NDRs elsewhere in the genome. In contrast, similarly sized random genomic regions display limited transcriptional activity. Moreover, Henriques et al. [55] recently found that, in S2 cells, enhancers with greater activity (as measured by a reporter assay [56]) are more highly transcribed (as measured by Start-seq). These findings further demonstrate the utility of high-resolution TSS profiling in elucidating potential functions of eRNAs.

Strikingly, we found that nuTSSs present within annotated enhancers are strongly unidirectional. In contrast, previous reports showed that, in S2 cells, putative eRNAs associated with computationally predicted intronic enhancers were significantly more divergent than active promoters [41]. Because the eRNAs we identified are associated with validated enhancer regions and are not enriched for downstream relative to upstream nuclear RNA-seq signal, we can be confident that our results are not significantly confounded by unannotated promoters. Therefore, either unidirectionality is a true characteristic of *D. melanogaster* eRNAs or perhaps the decay of one of the two presumptive eRNAs is much more rapid than the other. Nevertheless, the data point to a novel modality for eRNA genesis and function, perhaps distinct from the existing hypothesis that eRNAs originate from “underdeveloped” promoter regions that have not yet accumulated the cis-regulatory elements necessary to discriminate against antisense transcription [28].

## Materials and methods

### RNA library preparation and sequencing

For all libraries, nuclei were isolated from whole 3^rd^ instar *D. melanogaster* larvae as previously described [31]. For Nuclear RNA-seq and Start-seq, RNA was extracted from isolated nuclei using TRIzol reagent (Thermo Fisher). Start-seq libraries were prepared from nuclear RNA as previously described [19,21], and were sequenced on a NextSeq500 generating paired-end, 26nt reads. For nuclear RNA-seq, Total nuclear RNA was used as input to Ribo-zero Stranded RNA-seq library preparation (Illumina). Four biological replicates were prepared for Start-seq and nuclear RNA-seq. Libraries were sequenced on a HiSeq2000 generating paired-end, 50nt reads (Illumina).

### ATAC-seq library preparation and sequencing

ATAC-seq libraries were prepared as previously described [31]. For each replicate, nuclei from 10 whole 3rd instar larvae were isolated as per Start-seq and nuclear pellets were gently homogenized with wide-bore pipette tips in 50 uL ATAC-seq lysis buffer (10 mM Tris·Cl, pH 7.4, 10 mM NaCl, 3 mM MgCl^2^, 0.1% (v/v) Igepal CA-630). Homogenate was directly used as input to the Nextera DNA library preparation kit (NEB) for tagmentation of chromatinized DNA, as described in Buenrostro *et. al.* [30]. Three biological replicates were prepared. Libraries were sequenced on a HiSeq2000 generating single-end, 50nt reads (Illumina).

### Bioinformatic analysis

All raw data (fastq files) from ATAC-seq, Start-seq, and nuclear RNA-seq are available in the Gene Expression Omnibus (GEO) archive at NCBI under the following accession number: GSE96922. All ChIP-seq data were downloaded from modEncode (<u>www.modencode.org</u>). In all cases where possible, data were derived from the 3^rd^ instar larval time point as determined by modEncode developmental staging procedures. GEO accession numbers for modEncode data used in this study are as follows: H3K4me1: GSM1147329-32; H3K4me3: GSM1200083-86; H3K27ac: GSM1200071-74; H3K36me3: GSM1147189-92; H3: GSM1147289-92; BEAF32: GSM1256853-56. All histone PTM ChIP-seq data was normalized to H3 ChIP-seq data using the Deeptools bigwigCompare utility. Predicted nucleosome occupancy data was obtained from genome-wide nucleosome prediction tracks in *D. melanogaster* generated by the Eran Segal laboratory (<u>https://genie.weizmann.ac.il/software/nucleogenomes.html</u>). For ChIP-seq, ATAC-seq, and predicted nucleosome occupancy, metagene plots were generated using the Deeptools package [57]. All browser screenshots were captured from the UCSC Genome Browser [58].

### Start-seq peak assignment

Reads from start-seq FASTQ files were clipped to the first 26nt to remove adapter sequence, then mapped to the *D. melanogaster* dm3 genome build with Bowtie2 [59]. We used a custom script to quantify mapped Start-seq read density at individual nucleotides. Briefly, we parsed the SAM alignment ouput files from Bowtie2 by bitwise flag to select only first-in-pair reads (representing bona-fide initiation sites), then assigned the position (chromosome and nucleotide) of the first nucleotide in each read to a hash table, then combined the results from each replicate into a counts table. To normalize read depth, we first calculated the number of reads mapping to a set of spike-in control transcripts using the bedtools multicov utility [60]. For the number of raw reads (R) mapping to each spike-in transcript *t* \in *T* in each replicate *i* \in *n* (*R_ti_*), we normalized *R_ti_* to the geometric mean of all *{R_t1_,…,R_tn_}* to generate the normalized transcript value *N_ti_*. Finally, for each replicate we calculated a replicate normalization score *S_i_* by calculating the geometric mean of all *{N_1i_,…,N_Ti_}.* For initial analysis in Figs. 1 and 2, Start-seq reads were assigned to *D. melanogaster* TSSs defined from a previous study [19].

To generate Start-seq peak clusters likely to belong to the same TSS, we first calculated an FDR cutoff for bona-fide Start-seq peak detection at 9 normalized reads per nucleotide per biological replicate, based on sequencing depth. Then we clustered nucleotides meeting this threshold as follows: For each nucleotide *n_i_*, an edge was established with a neighboring nucleotide *n_j_* if it occurred within 5nt of *n_i_*. Then clusters were formed by including all nucleotides *n* that occurred between terminal nucleotides upstream (*n_u_*) and downstream (*n_d_*) that were bound to the cluster by only a single edge, and thus terminated the “chain.” For each cluster, the cluster “summit” was identified as the nucleotide containing the most mapped reads, and secondary and tertiary peaks were identified as containing the second- and third-most reads, respectively of any nucleotide in the cluster, if applicable. From the summit, we calculated the proportion of reads in the cluster contained within 2nt on either side of the summit.

To compare our peak clusters with those reported in Hoskins et al. and Nechaev et al., for each comparison we subsampled our data and the comparison data to 20 million or 5 million reads, respectively. We then generated peak clusters using the strategy outlined above, while varying the distance allowed to form an edge between two neighboring nucleotides (5, 10, 15, 25, 50, 75, or 100nt), and calculated the percentages of peaked, broad, and weak TSS clusters for each permutation as outlined in the text.

To assign Start-seq peak clusters to observed TSSs (obsTSSs) or novel TSSs (nuTSSs), we searched for overlap between our peak clusters and a list of coordinates for the first exons from every transcript in the dm3 gene annontation, and assigned those that overlapped as obsTSSs. All other clusters that did not map to a defined first exon were assigned as nuTSSs. Promoter regions defined by CAGE in *D. melanogaster* embryos were obtained from Hoskins *et al*. (2011), and “integrated promoters” that overlapped with obsTSS peaks were detected using the bedtools intersect function [60].

### Start-seq peak shape and pausing index

To calculate peak shape index (SI), we adapted a formula from Hoskins *et al*. [13]:

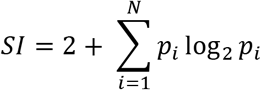

where *N* = the number of single nucleotides *i* in the peak cluster, and *p_i_* = the proportion of total reads in the cluster mapping to nucleotide *i*. To calculate peak pausing index (PI), we divided the normalized start-seq signal mapping to the obsTSS peak cluster by the nuclear RNA-seq RPKM calculated for its corresponding gene.

### Analysis of promoter motif enrichment

To discover motifs proximal to obsTSSs, we first sourced transcription factor motifs from FitzGerald *et al*. [38]. We then determined the expected distribution of those motifs relative to a TSS, and for every obsTSS we used the bedtools getfasta function to generate FASTA sequences that were 50nt in length and roughly restricted to the expected localization of each motif. For instance, the TATA box motif is expected to occur ~32nt upstream of the TSS, so the FASTA file used to test for the presence of TATA captured all nucleotides from -50 to 0 relative to the TSS. Then we used the 15 sequences present in FitzGerald *et al*. [38], plus the “Pause Button” motif sequence [34], and their corresponding restricted FASTA files for all obsTSSs, to execute the “homer2 find” function from the Homer motif analysis software [61], allowing for up to 4 mismatches. For each obsTSS *X* and each motif *y*, a single motif score *X_y_* was determined by selecting the highest log odds score among all *{X_y_1..X_y_n}* detected in the assigned FASTA sequence. Those obsTSSs for which a sequence with no more than 4 mismatches to a motif was not detected were assigned a score *X_y_* equal to the lowest log odds score for the all obsTSSs tested for the motif in question. *X_y_* values were converted to z-scores using the following formula:

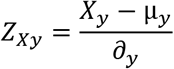

where *Z_X_y__* = z-score for obsTSS *X* and motif *y*, μ_y_ = mean log odds score for motif *y*, and *∂_y_* = standard deviation of log odds scores for motif *y*. The resultant z-scores were visualized using the heatmap.2 utility in the gplots R package (<u>https://CRAN.R-project.org/package=gplots</u>). with column clustering. Correlation coefficients for motifs were generated using the “cor” function in the R base package, and were visualized using the heatmap.2 utility.

### Divergent transcription analysis

To find instances of divergent transcription, pairs of neighboring TSSs oriented in opposite directions were ordered by distance of the reverse-strand TSS from the forward-strand TSS (negative numbers denote upstream, positive denote downstream), ATAC-seq signal was plotted in a 20kb window surrounding the forward-strand TSS using the Deeptools “computeMatrix reference-point” utility with default parameters [57], and signal was visualized using the heatmap.2 utility. From this analysis, divergent transcription was visually defined as the maximal distance between paired TSSs for which a conitiguous ATAC-seq enriched region could be detected, or roughly 200nt. For divergent pairs, Start-seq signal was visualized as described above, using computeMatrix reference-point with the following flags: -binSize 1, -afterRegionStartLength 1, -missingDataAsZero [57]. Enrichment of ATAC-seq signal around bidirectional, divergent, and non-divergent TSSs was determined by using the bedtools multicov tool [60] to map ATAC-seq signal within a 400nt window surrounding each TSS.

### nuTSS clustering and elongation analysis

To cluster nuTSSs by histone post-translational modification (PTM) density, PTM enrichment for H3K4me1, H3K4me3, H3K27ac, and H3K36me3 ChIP-seq signal at each nuTSS was calculated by mapping ChIP and input reads to a 200nt window on either side of the nuTSS using the bedtools multicov tool [60], then dividing ChIP reads by input reads. Optimal cluster number was discovered by calculating within-group sum of square distances for each cluster solution between 2 and 15 clusters, and 5 clusters were found to simultaneously minimize distance and cluster number. We assigned clusters using the hclust method in R.

To analyze transcription elongation from nuTSSs, we quantified the number of strand-specific nuclear RNA-seq reads mapping within either 200nt upstream or 200nt downstream of the nuTSS. Then, the enrichment of downstream over upstream reads was calculated using DESeq2 [62]. Though the expected paucity of reads mapping upstream of TSSs likely increases the threshold for significance across all TSSs, we reasoned that taking an approach that assigns significance partially based on the number of reads assigned to the feature in question would help in identify lowly elongating coding nuTSS transcripts oriented in the sense direction relative to their resident genes, since more reads would accumulate in both upstream and downstream regions in those cases.

### Enhancer region analysis

Functionally validated enhancers were obtained from the FlyLight database ([48], http://flweb.janelia.org/cgi-bin/flew.cgi) by querying based on anatomical expression in the larval CNS or in imaginal discs, and selecting all enhancers with validated expression in any one of those tissues. Each enhancer was annotated with the tissues in which it was reported to express according to the FlyLight database. Genomic coordinates of enhancer regions were obtained from the Bloomington Stock Center website (<u>http://flystocks.bio.indiana.edu/bloomhome.htm</u>). Meta-analysis of Start-seq signal probability in enhancer regions was conducted as follows: ATAC-seq peaks were called using Macs2 [63], and were classified based on overlap with gene TSSs, FlyLight enhancers, or neither by using consecutive instances of bedtools intersect [60]. Normalized Start-seq signal was mapped to a window of 2kb flanking either side of the peak summits using deeptools computeMatrix [57], and converted to binary (0 or 1) values to reflect presence or absence of signal. Separately, the same mapping and transformation was performed on a random permutation of the enhancer-overlapping set of peak summits. For each of the four sets, mean probability was calculated for each 10nt interval across the 4kb window, then each value normalized to the median of probability values within the set. Overlap between nuTSSs and enhancer regions was evaluated using the bedtools intersect tool [60].

## Declarations

### Ethics approval and consent to participate

Not applicable

### Consent for publication

Not applicable.

### Availability of data and materials

The datasets supporting the conclusions of this article are available in the Gene Expression Omnibus (GEO) repository, GSE96922 (https://www.ncbi.nlm.nih.gov/geo/query/acc.cgi?acc=GSE96922). Please see methods section for relevant GEO accession numbers for previously published modEncode data used in this manuscript.

### Author Contributions

MPM and AGM conceptualized the study and experiments. MPM designed methodology and performed formal analysis, validation, data curation, software design, and writing of the first draft. AGM, BDS, DJM, RJD and KA contributed to funding acquisition and review and editing.

### Acknowledgements

Not applicable.

---

**Supplementary Figure 1:**
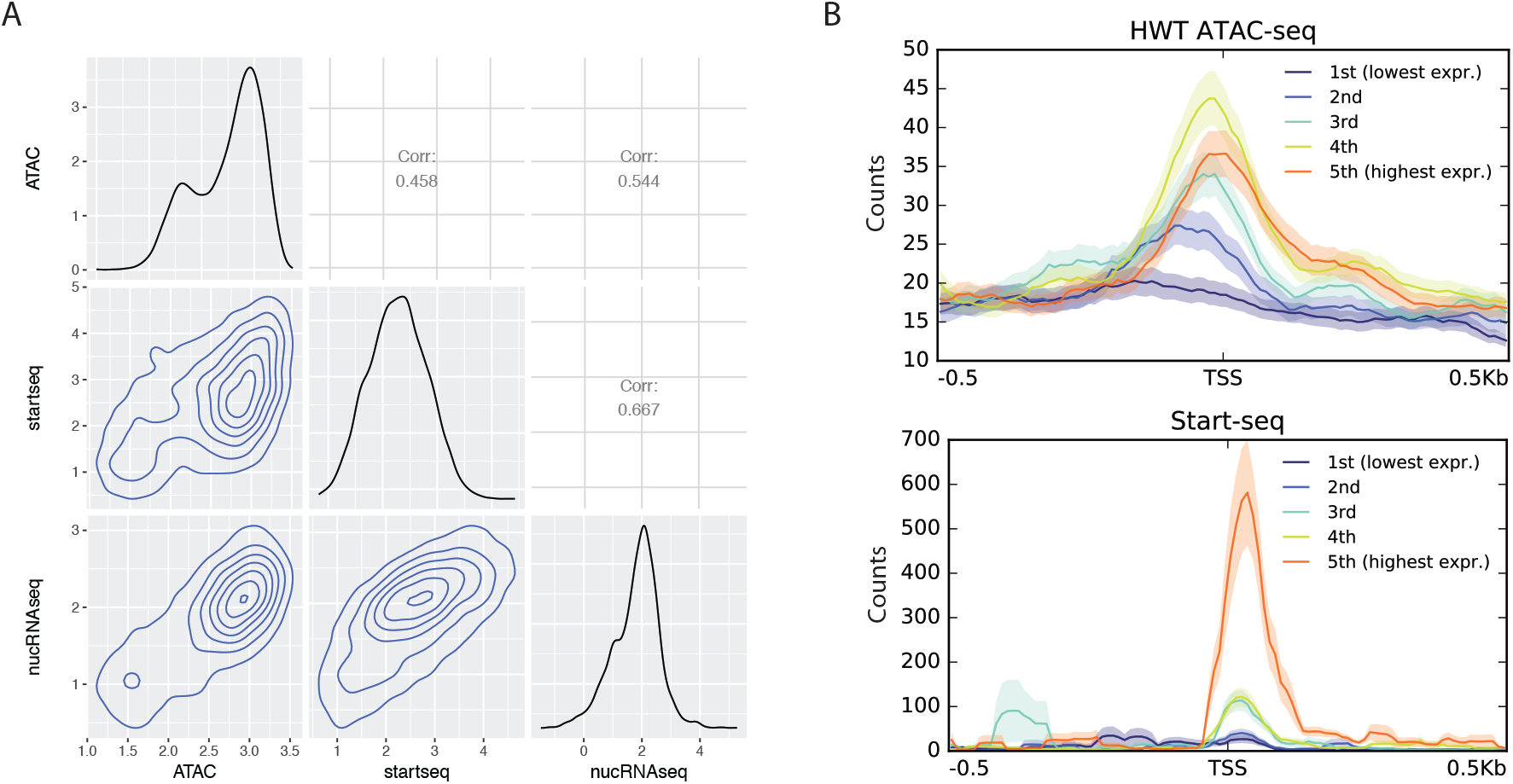
A) Pairwise comparisons of ATAC-seq, Start-seq, and nuclear RNA-seq for annotated coding genes. Boxes on diagonal show density plots of log signal enrichment for each assay. Boxes in on upper right show correlation values for each pairwise comparison. Boxes on lower left show contour plots representing scatterplots of log enrichment for each pairwise comparison. B) ATAC-seq and Start-seq signal accumulation at genes stratified into quintiles by nuclear RNA-seq signal. As is evident from lower ATAC-seq signal in the highest expression quintile, and by low stratification of lower quintiles by Start-seq, ATAC-seq and Start-seq are inconsistently correlated with nuclear RNA-seq.

**Supplementary Figure 2:**
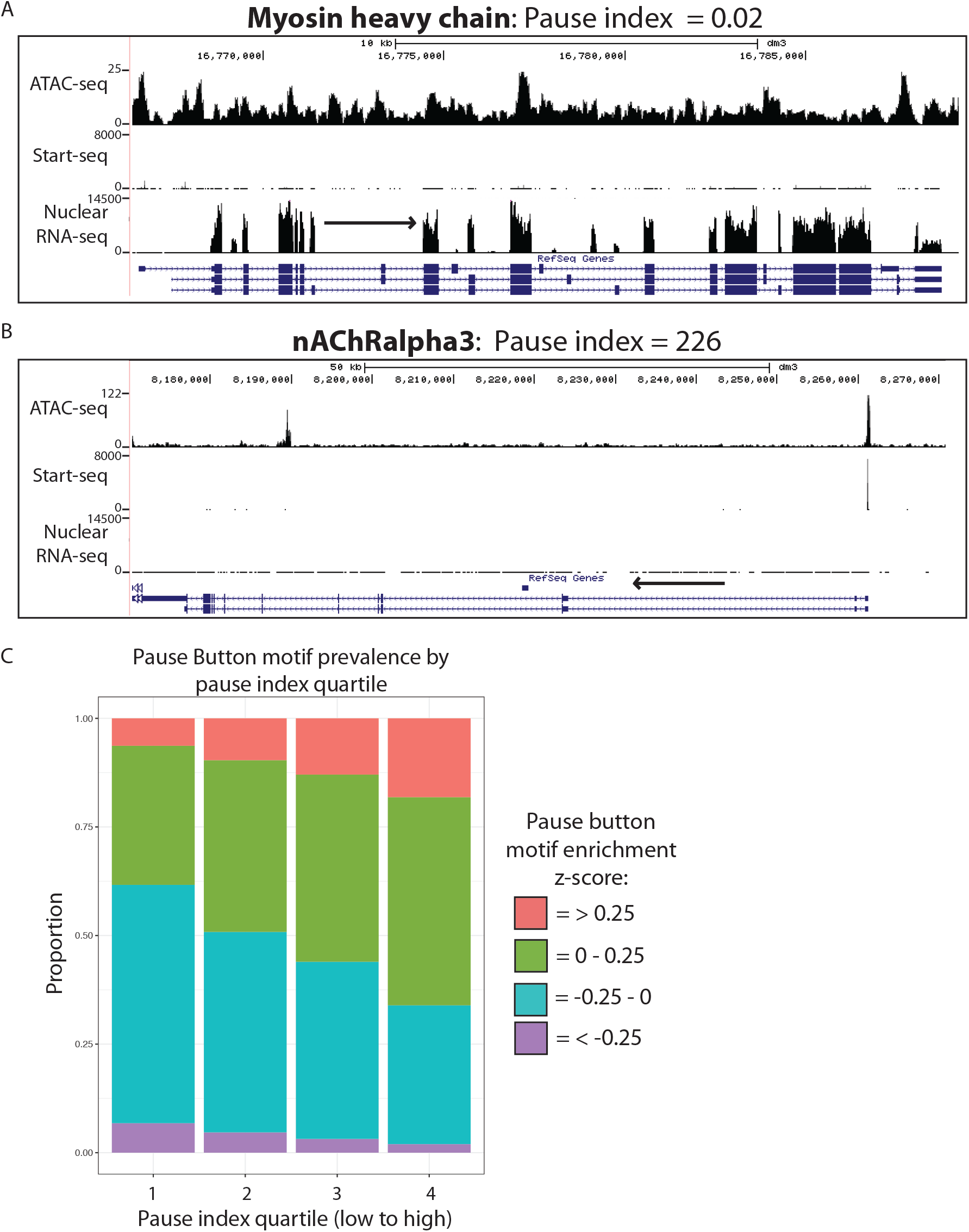
A) Representative browser window example of gene with low pause index. Note low Start-seq and high nuclear RNA-seq values. B) Representative browser window example of gene with high pause index. Note high Start-seq and low nuclear RNA-seq values. C) Enrichment of “Pause Button” (PB) motif based on pause index (PI) quartiles. Bars are partitioned based on z-score of highest log-odds score for enrichment of the PB motif in every nuTSS in the cohort. High PI quartile (right) disproportionately contains nuTSSs with high PB z-scores.

**Supplementary Figure 3:**
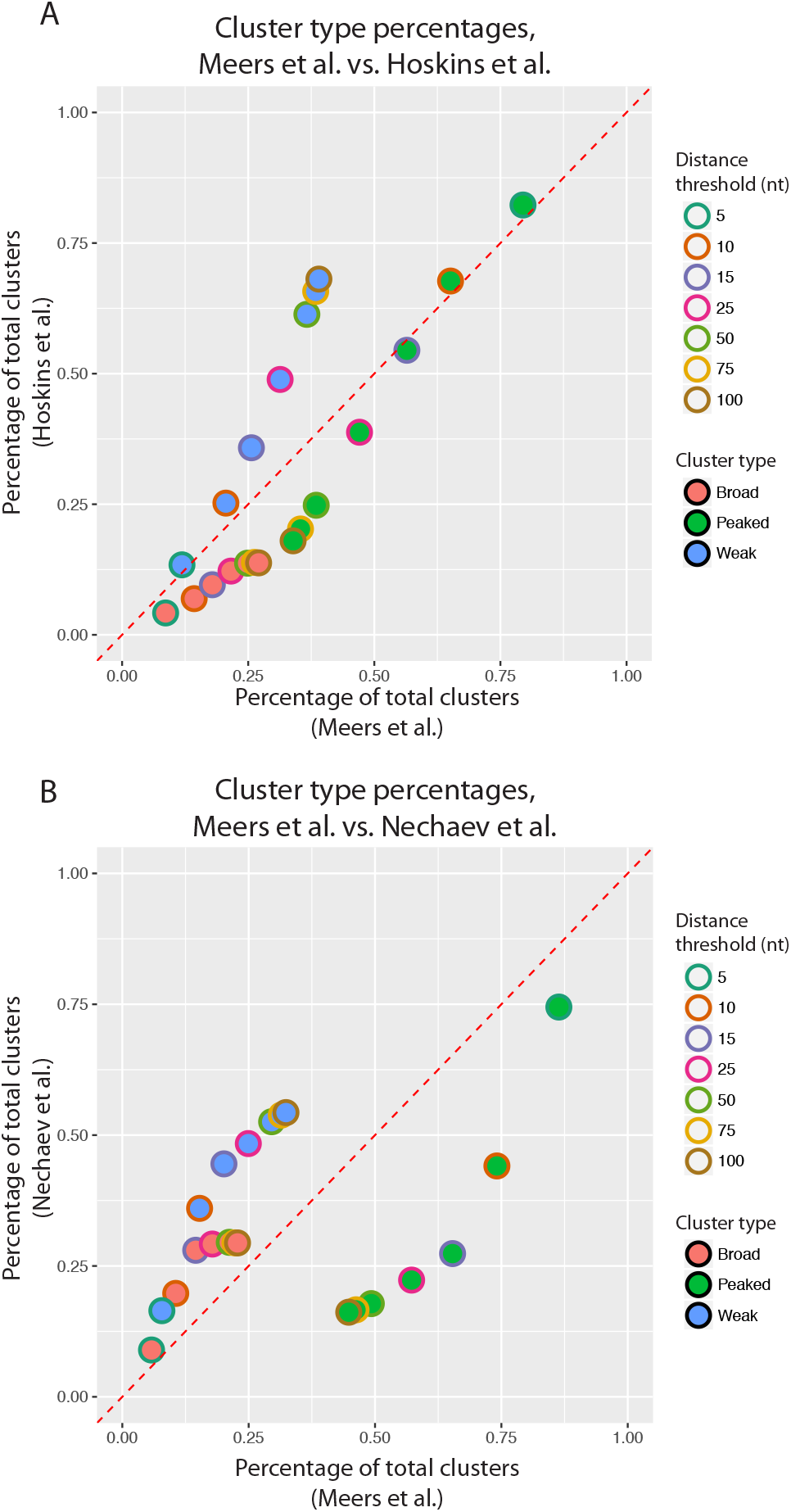
Comparison of the percentage of peaked (green points), broad (red), and weak (blue) clusters detected in this study (x-axis) as compared to Hoskins et al. (A) or Nechaev et al. (B) (y-axis). For each comparison, datasets were subsampled at equal read depth, signal was assigned to peaks using the same method, and the cluster distance threshold was varied (as denoted by colored point borders).

**Supplementary Figure 4:**
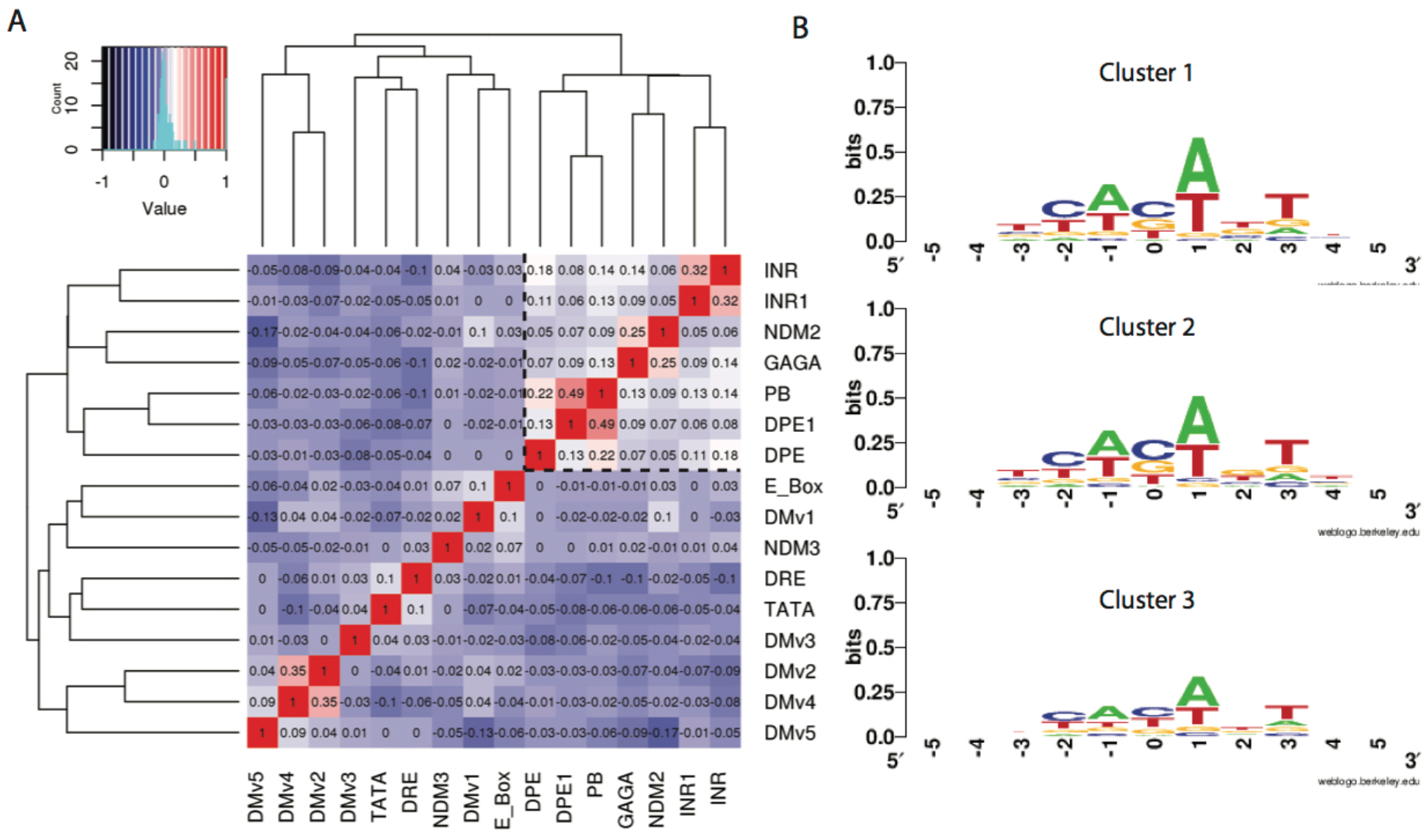
A) Correlation matrix of enrichment of 16 motifs used in clustering of obsTSSs. Dashed black lines denote group motifs co-occurring more frequently than others. B) Consensus motifs at obsTSSs based on motif-derived clustering. Cluster 1 resembles INR motif (TCAGT) from position -1 to 3, whereas other groups have lower sequence information content.

**Supplementary Figure 5:**
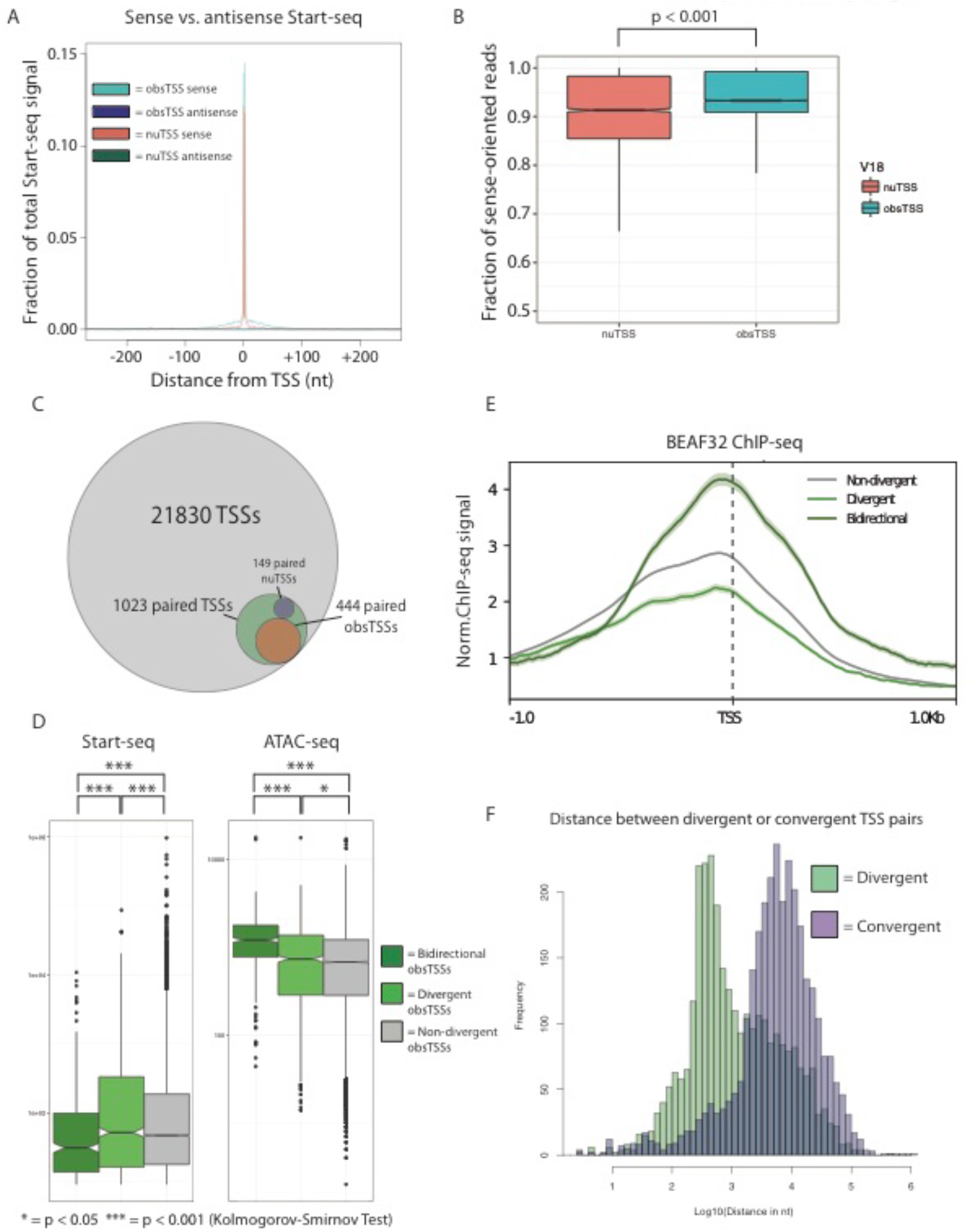
A) Metaplot of sense or antisense Start-seq signal mapping in a 500nt window around obsTSSs (teal and purple, respectively) or nuTSSs (salmon and green, respectively). B) Boxplot describing fraction of sense-oriented reads mapping in a 400nt window around nuTSSs (salmon) or obsTSSs (teal). C) Venn diagram describing representation of paired TSSs within high-confidence TSSs detected by Start-seq. D) Boxplots describing Start-seq (left) and ATAC-seq (right) signal corresponding to non-divergent obsTSSs (grey), divergent obsTSSs (i.e. obsTSSs paired with nuTSS, light green), and bidirectional obsTSSs (dark green). E) Metaplot describing BEAF32 ChlP-seq signal mapping in a 2kb window surrounding non-divergent (grey), divergent (light green), and bidirectional (dark green) obsTSSs. F) Histograms describing the distributions of distance between divergent (green) and convergent (purple) pairs of nearest neighbor obsTSSs oriented in opposite directions.

**Supplementary Figure 6:**
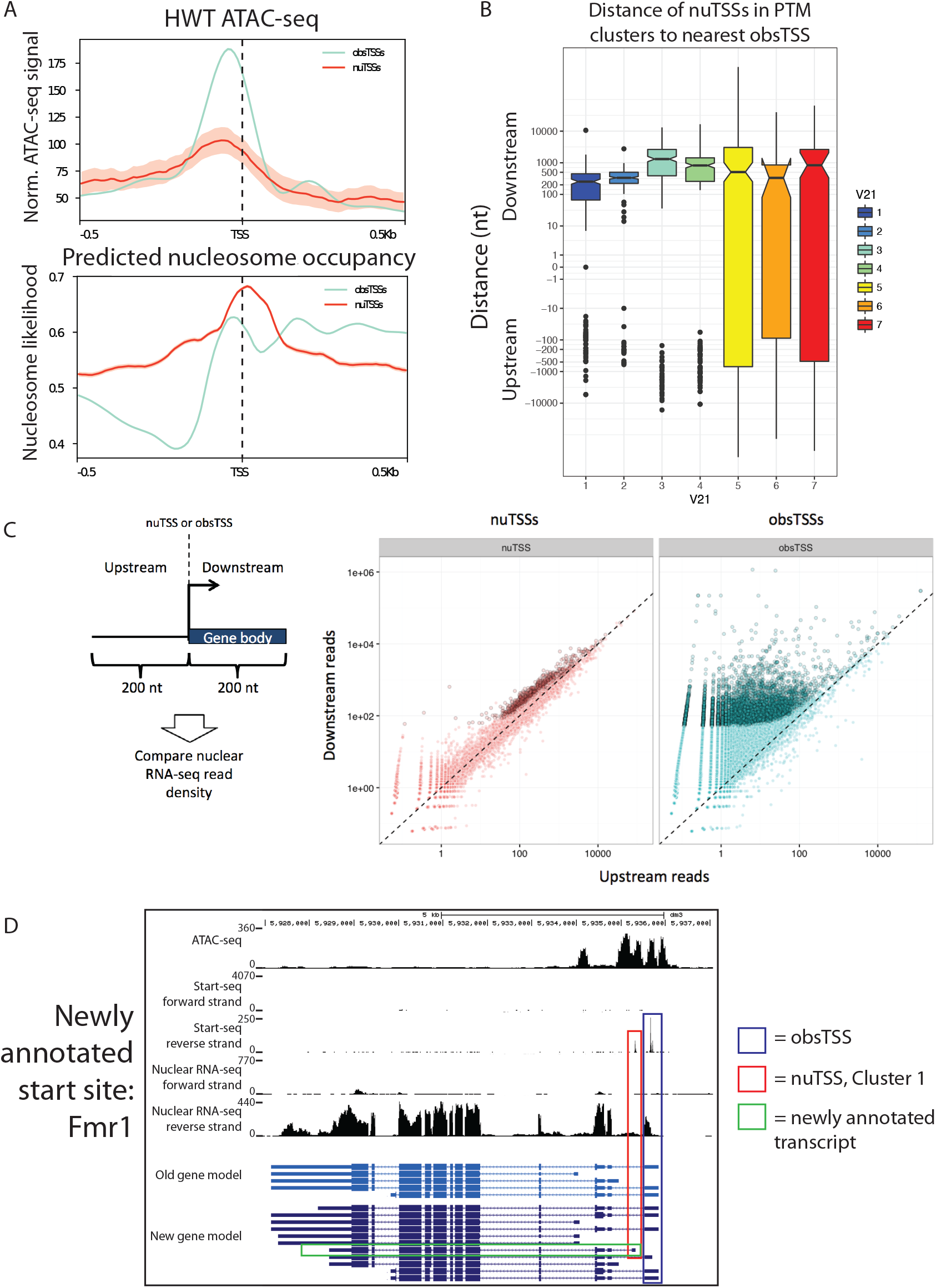
A) Metaplots describing ATAC-seq signal (top) and predicted nucleosome occupancy (bottom) in a 1 kb window surrounding obsTSSs (teal) or nuTSSs (red). B) Boxplot describing distribution of distances from the nearest obsTSS for all nuTSSs in each of 7 histone PTM-defined clusters described in Figure 5A. Negative values indicate nuTSSs that are upstream of the nearest obsTSS, while positive values indicate nuTSSs that are downstream of the nearest obsTSS. C) Nuclear RNA-seq reads were mapped to regions 200nt upstream and downstream of each TSS, and enrichment of downstream vs. upstream reads was analyzed as a proxy for elongation. At right: scatterplots of nuclear RNA-seq upstream reads (X-axis) vs. downstream reads (Y-axis) for nuTSSs (left, salmon) and obsTSSs (right, teal). D) Representative example of a nuTSS belonging to Cluster 1 (red box) that corresponds to a TSS that has been newly annotated in the latest *D. melanogaster* genome build (green box).

**Supplementary Figure 7:**
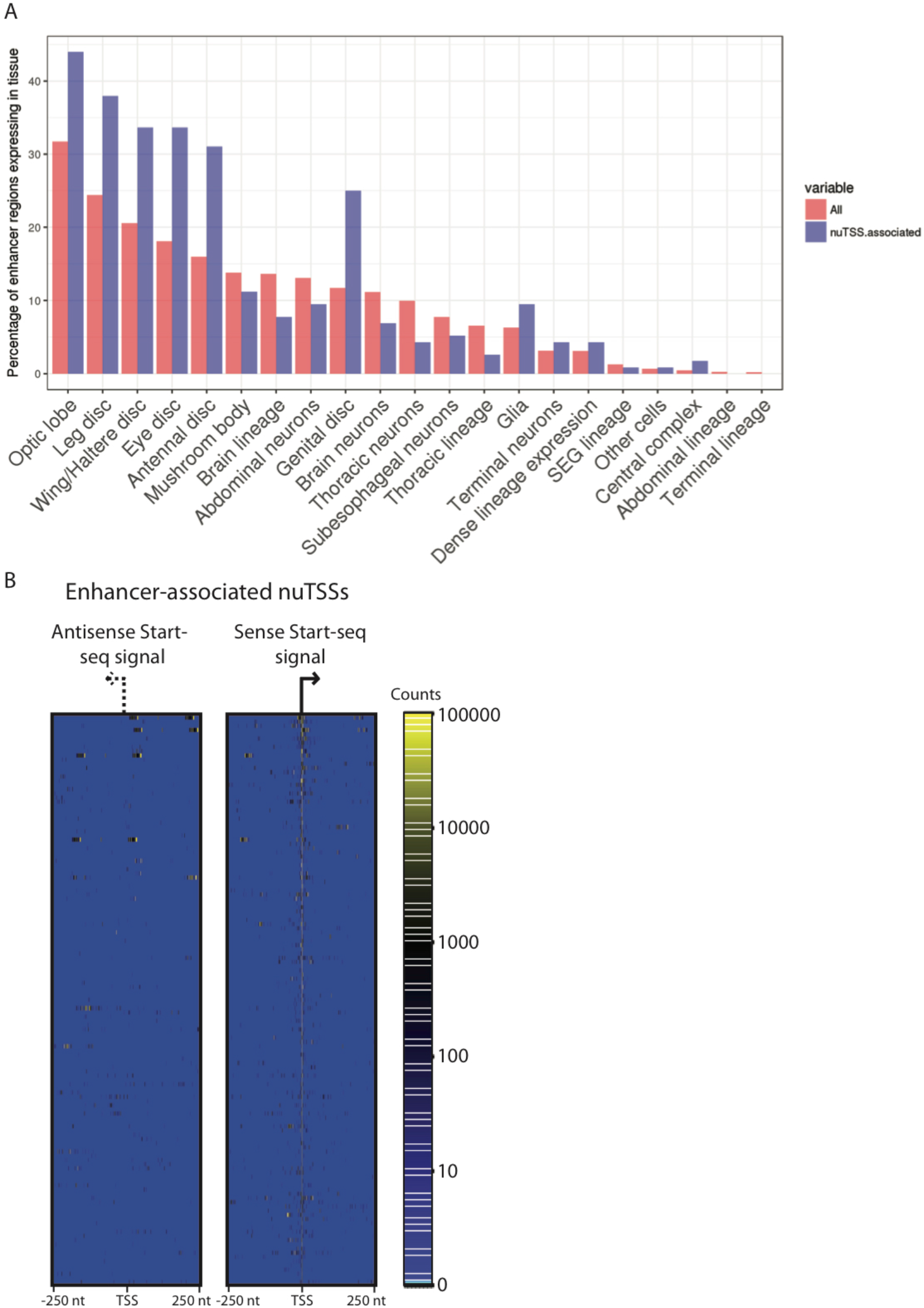
A) Percentage of all FlyLight enhancers (red) or those overlapping with a high-confidence nuTSS (blue) that correspond to each of several larval expression categories. B) Heatmap describing sense (right) or antisense (left) Start-seq signal mapping in a 500nt window around enhancer-associated nuTSSs.

## References

1. Buratowski S, Hahn S, Guarente L, Sharp PA. Five intermediate complexes in transcription initiation by RNA polymerase II. Cell 1989;56:549–61.

2. Mitchell PJ, Tjian R. Transcriptional regulation in mammalian cells by sequence-specific DNA binding proteins. Science 1989;245:371–8.

3. Lorch Y, Zhang M, Kornberg RD. Histone octamer transfer by a chromatin-remodeling complex. Cell 1999;96:389–92.

4. Segal E, Fondufe-Mittendorf Y, Chen L, Thåström A, Field Y, Moore IK, et al. A genomic code for nucleosome positioning. Nature 2006;442:772–8.

5. Kaplan N, Moore IK, Fondufe-Mittendorf Y, Gossett AJ, Tillo D, Field Y, et al. The DNA-encoded nucleosome organization of a eukaryotic genome. Nature. 2009;458:362–6.

6. Lee TI, Young RA. Transcription of eukaryotic protein-coding genes. Annu. Rev. Genet. 2000;34:77–137.

7. Carrozza MJ, Li B, Florens L, Suganuma T, Swanson SK, Lee KK, et al. Histone H3 methylation by Set2 directs deacetylation of coding regions by Rpd3S to suppress spurious intragenic transcription. Cell 2005;123:581–92.

8. Keogh M-C, Kurdistani SK, Morris SA, Ahn SH, Podolny V, Collins SR, et al. Cotranscriptional set2 methylation of histone H3 lysine 36 recruits a repressive Rpd3 complex. Cell 2005;123:593–605.

9. Takahashi K, Yamanaka S. Induction of pluripotent stem cells from mouse embryonic and adult fibroblast cultures by defined factors. Cell 2006;126:663–76.

10. Mavrich TN, Ioshikhes IP, Venters BJ, Jiang C, Tomsho LP, Qi J, et al. A barrier nucleosome model for statistical positioning of nucleosomes throughout the yeast genome. Genome Res. 2008;18:1073–83.

11. Ozsolak F, Song JS, Liu XS, Fisher DE. High-throughput mapping of the chromatin structure of human promoters. Nat. Biotechnol. 2007;25:244–8.

12. Carninci P, Sandelin A, Lenhard B, Katayama S, Shimokawa K, Pon_j_avic J, et al. Genome-wide analysis of mammalian promoter architecture and evolution. Nat. Genet. 2006;38:626–35.

13. Hoskins RA, Landolin JM, Brown JB, Sandler JE, Takahashi H, Lassmann T, et al. Genome-wide analysis of promoter architecture in Drosophila melanogaster. Genome Res. 2011;21:182–92.

14. Kim T-K, Hemberg M, Gray JM, Costa AM, Bear DM, Wu J, et al. Widespread transcription at neuronal activity-regulated enhancers. Nature 2010;465:182–7.

15. Djebali S, Davis CA, Merkel A, Dobin A, Lassmann T, Mortazavi A, et al. Landscape of transcription in human cells. Nature 2012;489:101–8.

16. Preker P, Nielsen J, Kammler S, Lykke-Andersen S, Christensen MS, Mapendano CK, et al. RNA exosome depletion reveals transcription upstream of active human promoters. Science 2008;322:1851–4.

17. Scruggs BS, Gilchrist DA, Nechaev S, Muse GW, Burkholder A, Fargo DC, et al. Bidirectional Transcription Arises from Two Distinct Hubs of Transcription Factor Binding and Active Chromatin. Mol. Cell 2015;58:1101–12.

18. Core LJ, Martins AL, Danko CG, Waters CT, Siepel A, Lis JT. Analysis of nascent RNA identifies a unified architecture of initiation regions at mammalian promoters and enhancers. Nat. Genet. 2014;46:1311–20.

19. Nechaev S, Fargo DC, dos Santos G, Liu L, Gao Y, Adelman K. Global analysis of short RNAs reveals widespread promoter-proximal stalling and arrest of Pol II in Drosophila. Science 2010;327:335–8.

20. Muse GW, Gilchrist DA, Nechaev S, Shah R, Parker JS, Grissom SF, et al. RNA polymerase is poised for activation across the genome. Nat. Genet. 2007;39:1507–11.

21. Henriques T, Gilchrist DA, Nechaev S, Bern M, Muse GW, Burkholder A, et al. Stable pausing by RNA polymerase II provides an opportunity to target and integrate regulatory signals. Mol. Cell 2013;52:517–28.

22. Saunders A, Core LJ, Sutcliffe C, Lis JT, Ashe HL. Extensive polymerase pausing during Drosophila axis patterning enables high-level and pliable transcription. Genes & Dev. 2013;27:1146–58.

23. Andersen PR, Domanski M, Kristiansen MS, Storvall H, Ntini E, Verheggen C, et al. The human cap-binding complex is functionally connected to the nuclear RNA exosome. Nat. Struct. Mol. Biol. 2013;20:1367–76.

24. Lubas M, Andersen PR, Schein A, Dziembowski A, Kudla G, Jensen TH. The human nuclear exosome targeting complex is loaded onto newly synthesized RNA to direct early ribonucleolysis. Cell Rep. 2015;10:178–92.

25. Lopez P, Wagner K-D, Hofman P, Van Obberghen E. RNA Activation of the Vascular Endothelial Growth Factor Gene (VEGF) Promoter by Double-Stranded RNA and Hypoxia: Role of Noncoding VEGF Promoter Transcripts. Mol. Cell. Biol. 2016;36:1480–93.

26. Kaplan CD, Laprade L, Winston F. Transcription elongation factors repress transcription initiation from cryptic sites. Science 2003;301:1096–9.

27. Verdel A, Jia S, Gerber S, Sugiyama T, Gygi S, Grewal SIS, et al. RNAi-mediated targeting of heterochromatin by the RITS complex. Science 2004;303:672–6.

28. Li W, Notani D, Rosenfeld MG. Enhancers as non-coding RNA transcription units: recent insights and future perspectives. Nat. Rev. Genet. 2016;17:207–23.

29. Vera JM, Dowell RD. Survey of cryptic unstable transcripts in yeast. BMC Genomics 2016;17:305.

30. Buenrostro JD, Giresi PG, Zaba LC, Chang HY, Greenleaf WJ. Transposition of native chromatin for fast and sensitive epigenomic profiling of open chromatin, DNA-binding proteins and nucleosome position. Nat. Methods 2013;10:1213–8.

31. Meers MP, Henriques T, Lavender CA, McKay DJ, Strahl BD, Duronio RJ, et al. Histone gene replacement reveals a post-transcriptional role for H3K36 in maintaining metazoan transcriptome fidelity. eLife 2017;6.

32. Gilchrist DA, Dos Santos G, Fargo DC, Xie B, Gao Y, Li L, et al. Pausing of RNA polymerase II disrupts DNA-specified nucleosome organization to enable precise gene regulation. Cell 2010;143:540–51.

33. Adelman K, Kennedy MA, Nechaev S, Gilchrist DA, Muse GW, Chinenov Y, et al. Immediate mediators of the inflammatory response are poised for gene activation through RNA polymerase II stalling. Proc. Natl. Acad. Sci. U.S.A. 2009;106:18207–12.

34. Hendrix DA, Hong J-W, Zeitlinger J, Rokhsar DS, Levine MS. Promoter elements associated with RNA Pol II stalling in the Drosophila embryo. Proc. Natl. Acad. Sci. U.S.A. 2008;105:7762–7.

35. Rach EA, Winter DR, Ben_j_amin AM, Corcoran DL, Ni T, Zhu J, et al. Transcription initiation patterns indicate divergent strategies for gene regulation at the chromatin level. PLoS Genet. 2011;7:e1001274.

36. Ni T, Corcoran DL, Rach EA, Song S, Spana EP, Gao Y, et al. A paired-end sequencing strategy to map the complex landscape of transcription initiation. Nat. Methods 2010;7:521–7.

37. Rye M, Sandve GK, Daub CO, Kawaji H, Carninci P, Forrest ARR, et al. Chromatin states reveal functional associations for globally defined transcription start sites in four human cell lines. BMC Genomics 2014;15:120.

38. FitzGerald PC, Sturgill D, Shyakhtenko A, Oliver B, Vinson C. Comparative genomics of Drosophila and human core promoters. Genome Biol. 2006;7:R53.

39. Trinklein ND, Aldred SF, Hartman SJ, Schroeder DI, Otillar RP, Myers RM. An abundance of bidirectional promoters in the human genome. Genome Res. 2004;14:62–6.

40. Yang J, Ramos E, Corces VG. The BEAF-32 insulator coordinates genome organization and function during the evolution of Drosophila species. Genome Res. 2012;22:2199–207.

41. Core LJ, Waterfall JJ, Gilchrist DA, Fargo DC, Kwak H, Adelman K, et al. Defining the status of RNA polymerase at promoters. Cell Rep. 2012;2:1025–35.

42. Mayer A, di Iulio J, Maleri S, Eser U, Vierstra J, Reynolds A, et al. Native elongating transcript sequencing reveals human transcriptional activity at nucleotide resolution. Cell 2015;161:541–54.

43. Liang G, Lin JCY, Wei V, Yoo C, Cheng JC, Nguyen CT, et al. Distinct localization of histone H3 acetylation and H3-K4 methylation to the transcription start sites in the human genome. Proc. Natl. Acad. Sci. U.S.A. 2004;101:7357–62.

44. Creyghton MP, Cheng AW, Welstead GG, Kooistra T, Carey BW, Steine EJ, et al. Histone H3K27ac separates active from poised enhancers and predicts developmental state. Proc. Natl. Acad. Sci. U.S.A. 2010;107:21931–6.

45. Rada-Iglesias A, Bajpai R, Swigut T, Brugmann SA, Flynn RA, Wysocka J. A unique chromatin signature uncovers early developmental enhancers in humans. Nature 2011;470:279–83.

46. Zentner GE, Tesar PJ, Scacheri PC. Epigenetic signatures distinguish multiple classes of enhancers with distinct cellular functions. Genome Res. 2011;21:1273–83.

47. Lipshitz HD, Peattie DA, Hogness DS. Novel transcripts from the Ultrabithorax domain of the bithorax complex. Genes & Dev. 1987;1:307–22.

48. Jenett A, Rubin GM, Ngo T-TB, Shepherd D, Murphy C, Dionne H, et al. A GAL4-driver line resource for Drosophila neurobiology. Cell Rep. 2012;2:991–1001.

49. Jory A, Estella C, Giorgianni MW, Slattery M, Laverty TR, Rubin GM, et al. A survey of 6,300 genomic fragments for cis-regulatory activity in the imaginal discs of Drosophila melanogaster. Cell Rep. 2012;2:1014–24.

50. McKay DJ, Lieb JD. A common set of DNA regulatory elements shapes Drosophila appendages. Dev. Cell 2013;27:306–18.

51. Dorighi KM, Swigut T, Henriques T, Bhanu NV, Scruggs BS, Nady N, et al. Mll3 and Mll4 Facilitate Enhancer RNA Synthesis and Transcription from Promoters Independently of H3K4 Monomethylation. Mol. Cell 2017;66:568–576.e4.

52. Wakano C, Byun JS, Di L-J, Gardner K. The dual lives of bidirectional promoters. Biochim. Biophys. Acta. 2012;1819:688–93.

53. Duttke SHC, Lacadie SA, Ibrahim MM, Glass CK, Corcoran DL, Benner C, et al. Human promoters are intrinsically directional. Mol. Cell 2015;57:674–84.

54. Hallikas O, Palin K, Sin_j_ushina N, Rautiainen R, Partanen J, Ukkonen E, et al. Genome-wide prediction of mammalian enhancers based on analysis of transcription-factor binding affinity. Cell 2006;124:47–59.

55. Henriques T, Scruggs BS, Inouye MO, Muse GW, Williams L, Burkholder AB, Lavender CA, Fargo DC and Adelman K. Widespread transcriptional pausing and elongation control at enhancers. Genes & Dev. (in press).

56. Arnold CD, Gerlach D, Stelzer C, Boryń ŁM, Rath M, Stark A. Genome-wide quantitative enhancer activity maps identified by STARR-seq. Science 2013; 339:1074–1077.

57. Ramírez F, Dündar F, Diehl S, Grüning BA, Manke T. deepTools: a flexible platform for exploring deep-sequencing data. Nucleic Acids Res. 2014;42:187–191.

58. Kent WJ, Sugnet CW, Furey TS, Roskin KM, Pringle TH, Zahler AM, et al. The human genome browser at UCSC. Genome Res. 2002;12:996–1006.

59. Langmead B, Salzberg SL. Fast gapped-read alignment with Bowtie 2. Nat. Methods 2012;9:357–9.

60. Quinlan AR, Hall IM. BEDTools: a flexible suite of utilities for comparing genomic features. Bioinforma. Oxf. Engl. 2010;26:841–2.

61. Heinz S, Benner C, Spann N, Bertolino E, Lin YC, Laslo P, et al. Simple combinations of lineage-determining transcription factors prime cis-regulatory elements required for macrophage and B cell identities. Mol. Cell 2010;38:576–89.

62. Love MI, Huber W, Anders S. Moderated estimation of fold change and dispersion for RNA-seq data with DESeq2. Genome Biol. 2014;15:550.

63. Zhang Y, Liu T, Meyer CA, Eeckhoute J, Johnson DS, Bernstein BE, et al. Model-based analysis of ChIP-Seq (MACS). Genome Biol. 2008;9:R137.

